# CD160 is an inducible endothelial checkpoint that suppresses sprouting angiogenesis through LRP1-dependent autophagy

**DOI:** 10.64898/2026.10.19.760050

**Authors:** Abdel-ilah Aziz, Maxence Mocquery-Corre, Alexandre Berquand, Jessica Thevenard-Devy, André Martins, Cathy Hachet, Stéphane Potteaux, Yacine Merrouche, Rachid El Fatimy, Laurent Duca, Armand Bensussan, Sanae Ben Mkaddem, Jérôme Devy

## Abstract

Pathological angiogenesis is a hallmark of cancer and other chronic diseases, yet therapeutic strategies remain largely focused on blocking pro-angiogenic signals rather than restoring endogenous mechanisms that restrain endothelial sprouting. The identification of endothelial checkpoints capable of limiting pathological vessel growth therefore remains an important unmet need. CD160 is a glycosylphosphatidylinositol-anchored immunoglobulin-superfamily receptor of cytotoxic lymphocytes and an inhibitory checkpoint receptor on CD4 T cells. It is also present on endothelium and has been targeted therapeutically. Here, we identify CD160 as an inducible endothelial checkpoint that emerges in response to pro-angiogenic cues and limits pathological angiogenesis. CD160 is absent in resting endothelium. It is induced by basic fibroblast growth factor (bFGF) and most strongly by combining VEGF-A and bFGF, and it is detected in the sprouting region of *ex vivo* mouse aortic rings. Targeting of CD160 using monoclonal antibody reduces tubulogenesis, scratch closure and aortic-ring sprouting, and softens the cell cortex, while viability and membrane integrity remain intact. Mechanistically, CD160 associates with the endocytic receptor LRP1 as demonstrated by proximity ligation assay, lipid-raft co-immunoprecipitation and super-resolution fluorescence microscopy. Both receptors appear in a common complex, and two independent computational methods converge on the first ligand-binding cluster of LRP1 as the interface. Silencing CD160 or LRP1 abolishes the anti-angiogenic response mediated by the antibody. Under a chloroquine clamp, engagement raises autophagic flux in a CD160- and LRP1-dependent manner, and lysosomal or LDL-receptor-family blockade removes the anti-sprouting effect. CD160 therefore acts as an inducible endothelial checkpoint that suppresses sprouting through autophagic flux. Thus, CD160 represents an inducible endothelial checkpoint that is activated in a pro-angiogenic environment, and targeting CD160 restrains pathological angiogensis.

## Introduction

Approved anti-angiogenic therapies primarily target the pathological activation of pro-angiogenic signaling pathways, through VEGF-A neutralization using monoclonal antibodies or inhibition of VEGF receptor kinase activity using small molecules^1,2^. Namely, anti-VEGF-A antibodies such as bevacizumab, the VEGFR-2 antibody ramucirumab and the ligand trap aflibercept^1^, together with receptor tyrosine kinase inhibitors such as sunitinib, sorafenib, axitinib and lenvatinib^2^. Their clinical benefit is usually transient. Tumours escape VEGF blockade by up-regulating alternative pro-angiogenic factors, by recruiting pro-angiogenic myeloid cells and by co-opting existing vessels, and a proportion of tumours are indifferent to the blockade from the outset^3^. This is a further argument for acting on the endothelium itself rather than on the ligand alone. However, vascular homeostasis is not governed solely by the suppression of pro-angiogenic cues; it also relies on endogenous inhibitory mechanisms that actively constrain endothelial activation and sprouting. This raises the possibility of therapeutically restoring these intrinsic vascular checkpoints to counteract pathological angiogenesis. Three paradigms support this approach. Netrin-1 engagement of UNC5B inhibits endothelial sprouting and promotes the retraction of endothelial filopodia^4,5^. DLL4–Notch1 signaling limits tip-cell selection and thereby limits sproutig^6^. Agonism of the angiopoietin–TIE axis stabilizes the vessel wall^7^. Each of these receptors is expressed or induced in activated endothelium, so engaging them is an agonist strategy that acts only where the vessel is already activated.

We refer to this class of receptors as endothelial checkpoint : inducible cell surface receptor that actively constrain endothelial activation and limit the angiogenic response. This concept extends the immune checkpoint paradigm to vascular pathophysiology, where engagement of an inducible inhibitory receptor limits an activated effector program^8^. The analogy is particularly relevant to CD160, an inhibitory checkpoint receptor expressed by human CD4 T cells, where it signals through herpesvirus entry mediator ^9^.

CD160 was identified as an 80-kDa cell-surface structure on circulating lymphocytes with natural-killer activity^10^ and was defined by the BY55 monoclonal antibody^11^. Cloning identified a 27-kDa immunoglobulin-superfamily protein with a single Ig-like domain, attached to the membrane by a glycosylphosphatidylinositol (GPI) anchor^12^. The 80-kDa form is a tightly disulfide-linked multimer of this 27-kDa monomer: reduction alone yields the 80-kDa form, whereas reduction combined with carboxyamidomethylation yields the 27-kDa monomer^12^. A transmembrane isoform was later described on activated NK cells^13^. Three forms are now established: GPI-anchored^12,14^, transmembrane^13^ and soluble^15^.

Previous work has placed CD160 in the regulation of vascular growth and remodeling. Soluble HLA-G1 binds CD160 on endothelial cells and inhibits tube formation, an effect associated with the induction of an apoptotic pathway^16^. Consistet with a anti-angiogenic function, CD160-targeting antibody inhibits angiogenesis *in vivo* and promotes tumor vessels normalization ^17^, and a same approach inhibits ocular neovascularization in rabbit and monkey models, alone or combined with bevacizumab^18^. Despite these therapeutic observations, the underlying mechanism remains unresolved. In particular, the proposed model in which CD160 engagement suppresses angiogenesis primarily through endothelial apoptosis does not establish whether cell death is the initiating event or a downstream consequence of altered endothelial function. However, In previous study the functional readout preceded the death readout^16^, so cell death can not be directly linked to the loss of angiogenic phenotype.

Furthermore, CD160 ligand; HLA-G engages several receptors. HLA-G operates as an immune checkpoint molecule through a defined receptor repertoire^19^. Its soluble form circulates as a disulfide-linked dimer that signals efficiently through the leukocyte immunoglobulin-like receptors LILRB1 and LILRB2^20^, through KIR2DL4 from endosomes^21^ and through CD8, where it triggers CD95/CD95L-mediated apoptosis in activated CD8+ cells^22^. These are leukocyte receptors, and none has been reported on endothelial cells. In tissue the situation differs. Soluble HLA-G1 reaches the leukocyte receptors as well, endothelial cells themselves express CD95L and use it to regulate leukocyte extravasation^23^ and to form an immune barrier in tumor vessels^24^, and a vascular outcome obtained with the HLA-G1 need not be endothelium-autonomous. Engagement with a CD160-specific agonist antibody, combined with CD160 silencing, isolates the CD160 contribution, and that is the approach used here.

The most important question that is still not answered two decades after the receptor was found on endothelium, its regulation and the molecule that carries its signal into the cell remain to be defined. As a GPI-anchored receptor is deprived of cytoplasmic tail, its signaling generally depends on a transmembrane partner^25^. Engagement of CD160 on circulating human NK cells drives its association with CD2^26^. In endothelium no partner has been yet reported. In this context, LRP1 appears as a candidate. It is a large endocytic receptor whose cytoplasmic tail binds numerous adaptors through NPxY, YxxL and LL sequences^27^ and which modulates other transmembrane receptors^28^. Two GPI-anchored receptors depend on it to operate. The urokinase plasminogen activator receptor (uPAR) requires direct binding to LRP1 for its endocytosis and for the surface urokinase activity it controls^29^, and the cellular prion protein (PrP^c^) uses LRP1 as its endocytic adaptor for clathrin^30^. LRP1 is active in endothelium, where it regulates retinal angiogenesis and endothelial proliferation^31,32^ and contributes to tumor angiogenesis^33,34^; antagonizing the uPAR–LRP1 interaction suppresses angiogenesis in an organotypic human iris model^35^. LRP1 also link ligand engagement to autophagy^36^, and its loss lowers autophagic flux in differentiating chondrocytes, where LC3-II and autophagosome number fall and p62 accumulates even though the upstream initiators Beclin-1, Fis1 and Perk are raised^37^.

This leads us to autophagy as a more plausible mechanism for CD160 engagement. Endothelial autophagy is required for sprouting: it shapes tumor vessel formation^38^, supports sprouting from aortic endothelium^39^, and endothelial deletion of *Atg5* attenuates ischemia-driven angiogenesis^40^. The pathway is controlled in both directions. Endothelial TFEB promotes post-ischemic angiogenesis through lysosomal biogenesis^41^, endothelial protein kinase A restrains angiogenesis by limiting autophagy through ATG16L1^42^, and lysosomal and autophagic routes have separable effects on endothelial behavior^43^.

Induced autophagy is itself angiostatic, and this is where established angiogenesis inhibitors converge. Kringle 5 of plasminogen induces autophagy in endothelial cells^44^; endostatin induces it by raising Beclin-1 and lowering β-catenin^45^; autophagy induction is a shared element of these inhibitors^46^. Some of matrix-derived angiostatic proteins act the same way. Decorin evokes endothelial autophagy through Peg3^47^, and endorepellin-evoked autophagy contributes to angiostasis^48,49^. Autophagy also degrades endothelial VEGFA, which lowers the autocrine drive for sprouting^50^. A receptor that raises endothelial autophagic flux is therefore expected to suppress sprouting.

Here we define CD160 as a growth-factor-inducible endothelial receptor, identify LRP1 as its membrane partner, and show that engaging CD160 suppresses sprouting while raising autophagic flux. Both responses require both receptors. Engagement leaves viability intact and softens the cortex, so the suppressed state is a viable, mechanically altered endothelium. Autophagy is measured as flux under a lysosomal clamp^51,52^, tube formation is used as a screening assay, and the *ex vivo* aortic ring provides the multicellular sprouting readout^53^. These findings identify a previously unrecognized endogenous brake on angiogenesis and implicate LRP1-dependent autophagic flux as an effector pathway through which endothelial CD160 limits pathological sprouting.

## Results

### CD160 is induced in human endothelium by bFGF and by VEGF-A + bFGF

An endothelial checkpoint should appear with the activated state. We stimulated HUVEC for 24 h with bFGF (20 ng mL−1), VEGF-A (50 ng mL−1), VEGF-A plus bFGF, TNF-α (10 ng mL−1) or TGF-β (5 ng mL−1). Background-corrected CD160 and VE-cadherin fluorescence were then measured per cell by confocal microscopy. Each marker was quantified inside a cell mask derived from the other (Fig. 1a,b). CD160 was at the detection floor in unstimulated cells (≈ 4 mean fluorescence intensity per cell). bFGF raised it to ≈ 43 (P = 0.0092 versus unstimulated) and VEGF-A plus bFGF to ≈ 61 (P = 0.0004). VEGF-A alone gave an intermediate signal that did not reach significance against the unstimulated arm (≈ 20). Neither TNF-α nor TGF-β raised CD160 above baseline (≈ 4 and ≈ 5). The combination exceeded VEGF-A alone (P = 0.0056). Increments over baseline were approximately additive (bFGF ≈ +39, VEGF-A ≈ +16, combination ≈ +57), so the two growth factors contribute independently. CD160 staining was membrane-associated in every positive condition (Fig. 1a). Membrane VE-cadherin was unchanged across the same six conditions (one-way ANOVA P = 0.29; Fig. 1b, right), so the CD160 response is neither a general consequence of endothelial activation nor an artefact of altered antigen accessibility.

**Figure 1.**
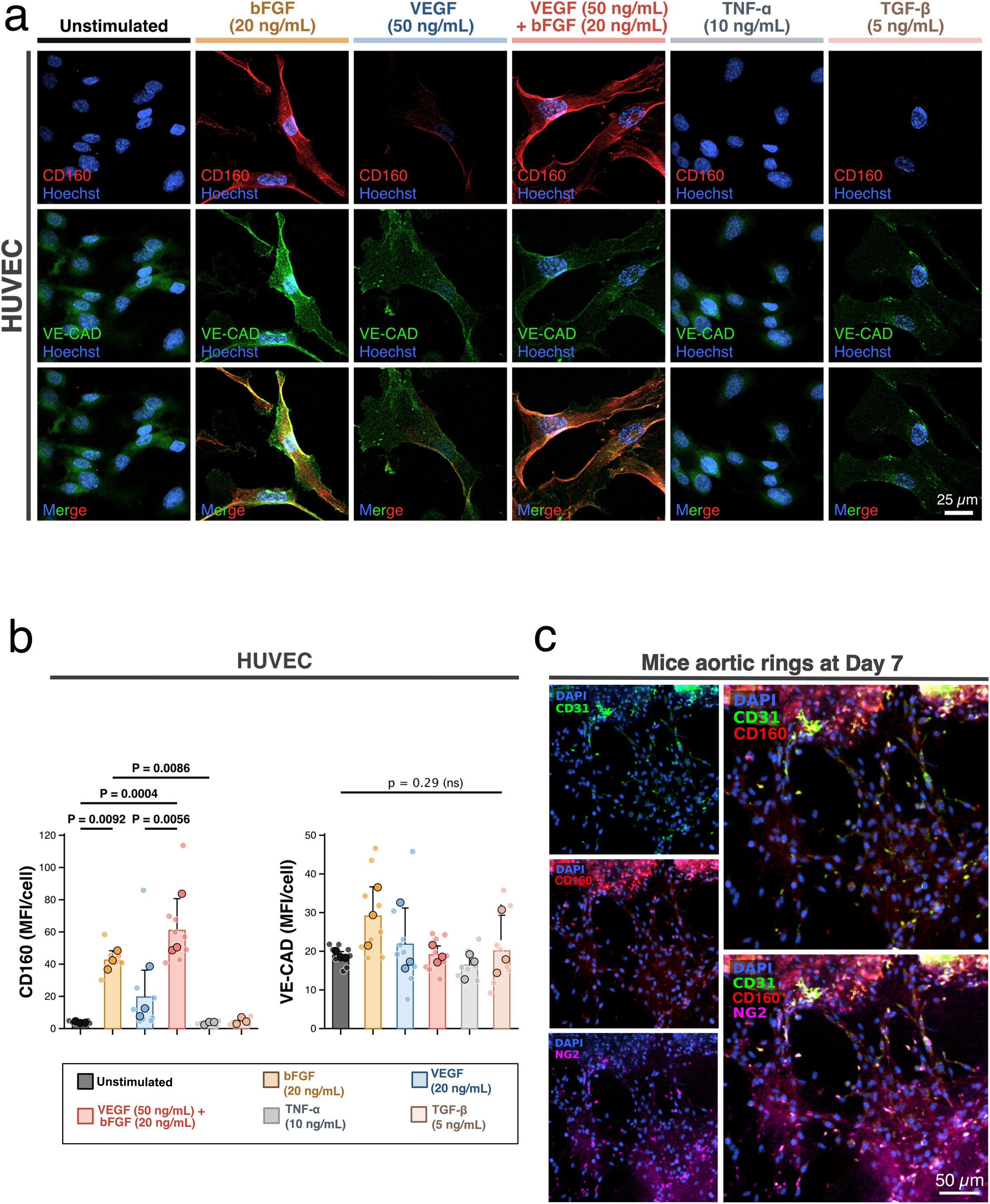
CD160 is induced in human endothelium by bFGF and by VEGF-A plus bFGF, and is present in sprouting aorta. **a**, Confocal immunofluorescence of HUVEC after 24 h in defined medium alone (unstimulated) or with bFGF (20 ng mL^−1^), VEGF-A (50 ng mL^−1^), VEGF-A plus bFGF, TNF-α (10 ng mL^−1^) or TGF-β (5 ng mL^−1^). Rows: CD160 (red) with Hoechst (blue); VE-cadherin (green) with Hoechst; merge. Acquisition settings identical across all conditions. Scale bar, 25 µm. **b**, Quantification of **a** as background-corrected mean fluorescence intensity per cell. Left, CD160; right, VE-cadherin. Each marker was quantified inside a cell mask derived from the other marker. One-way ANOVA with Dunnett’s multiple-comparison test against the unstimulated arm for the induction comparisons, with the additional planned comparisons of the combination against each single factor. CD160: *P* = 0.0092 (bFGF versus unstimulated), *P* = 0.0004 (VEGF-A plus bFGF versus unstimulated), *P* = 0.0056 (combination versus bFGF) and *P* = 0.0086 (combination versus VEGF-A); VEGF-A, TNF-α and TGF-β versus unstimulated were not significant. VE-cadherin: overall ANOVA *P* = 0.29, not significant. *n* = 3 independent experiments. **c**, Immunofluorescence of mouse aortic-ring outgrowths at day 7. Left column, single channels: DAPI with CD31 (green); DAPI with CD160 (red); DAPI with NG2 (magenta). Right column, merges of DAPI/CD31/CD160 and DAPI/CD31/CD160/NG2. Representative of 8 animals. Scale bar, 50 µm. This panel is qualitative; no quantification is presented.

The growth-factor dependence is specific. bFGF is the effective single inducer, VEGF-A alone is not sufficient, the two together are the strongest condition, and two canonical activating cytokines do not induce the receptor. CD160 is therefore inducible rather than constitutive in human endothelium.

CD160 was also present in a sprouting vessel wall. Day-7 mouse aortic-ring outgrowths were stained for CD31, CD160 and the mural-cell marker NG2 (Fig. 1c). CD160 immunoreactivity was detected in the sprouting region together with CD31, with NG2 marking a distinct mural compartment.

Antibody performance was characterized separately (Supplementary Fig. 1a). Anti-CD160 and the reference BY55 monoclonal^11^ both stained the CD160-positive YT lymphocyte line and CHO cells stably expressing CD160-GPI, and neither stained mock-transfected CHO cells. Staining of non-permeabilized cells places the epitope in the CD160 ectodomain.

### Engaging CD160 suppresses endothelial network formation without cytotoxicity

We investigated whether engaging CD160 changes angiogenic behavior. Anti-CD160 was compared with isotype-matched IgG1 in the HUVEC tube-formation assay, in growth-factor-supplemented EGM-2 (Fig. 2a,b). Anti-CD160 reduced total tube length in a dose-dependent manner. The difference reached significance at 10 and 20 µg mL−1 (≈ −39%, P = 0.023; ≈ −40%, P = 0.046) and not at 1 or 5 µg mL−1 (≈ −17%, P = 0.182; ≈ −21%, P = 0.055). Junction counts behaved the same way, with significance at 10 µg mL−1 (P = 0.047) and a consistent downward trend elsewhere (P = 0.223, 0.089 and 0.073 at 1, 5 and 20 µg mL−1; Fig. 2b, right). Expanded morphometry across the same dose series gives a steeper dose slope for anti-CD160 than for IgG1 in network area (−3.66 versus −1.72 per log10 unit), node number (−102.3 versus −71.6) and segment number (−42.8 versus −33.7); mesh number was the one parameter in which the IgG1 slope was steeper (Supplementary Fig. 2a). All subsequent experiments used 10 µg mL−1.

**Figure 2.**
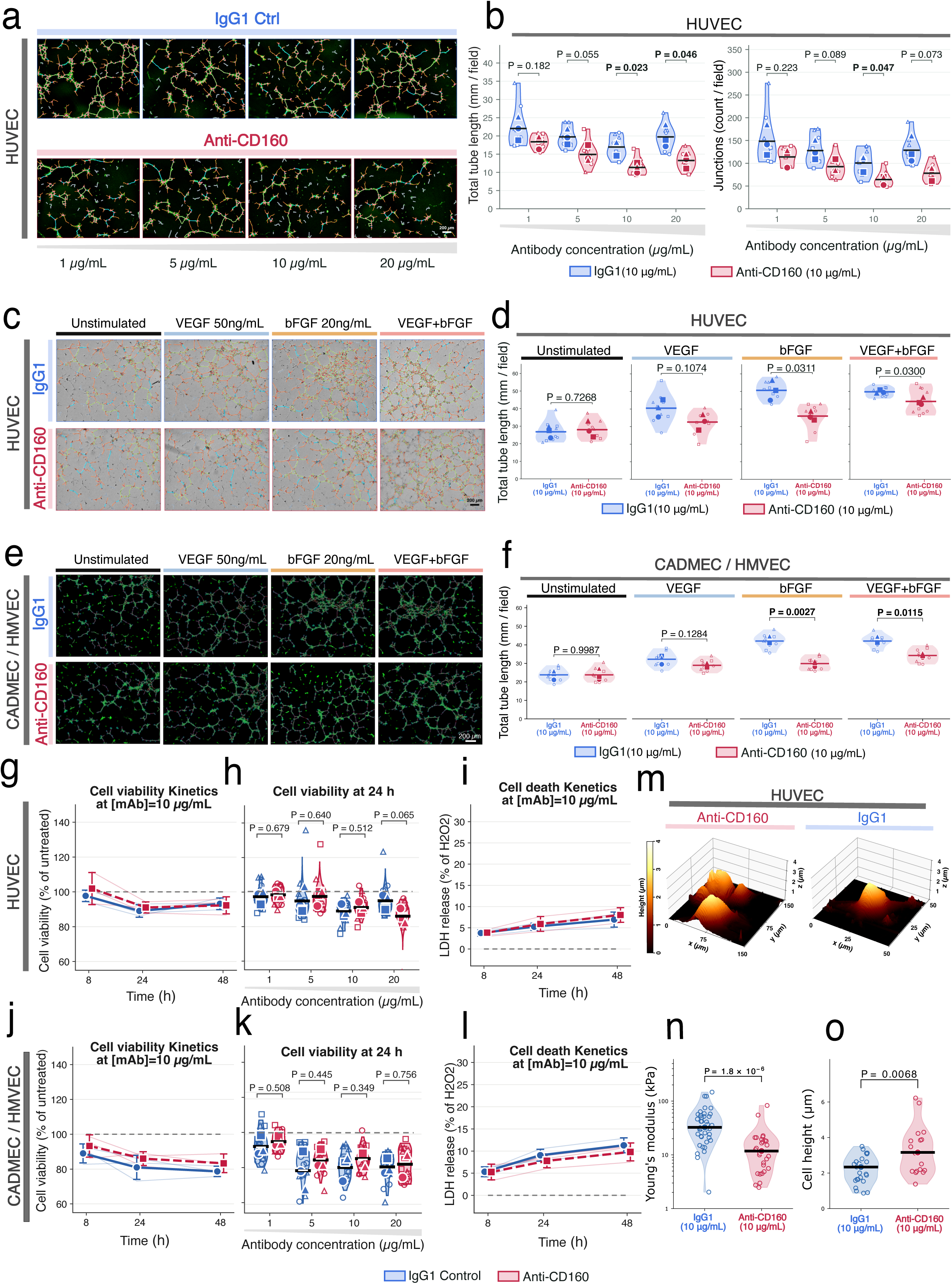
Antibody engagement of CD160 suppresses endothelial network formation without cytotoxicity and softens the cell cortex. **a**, HUVEC tube-formation networks after 6 h on growth-factor-reduced Matrigel with IgG1 (top) or anti-CD160 (bottom) at 1, 5, 10 and 20 µg mL^−1^, shown with the Angiogenesis Analyzer skeleton and branch-point overlay. Scale bar, 200 µm. **b**, Quantification of **a**. Left, total tube length (mm per field): *P* = 0.182, 0.055, 0.023 and 0.046 at 1, 5, 10 and 20 µg mL^−1^. Right, junction count per field: *P* = 0.223, 0.089, 0.047 and 0.073. Welch’s *t*-test, IgG1 versus anti-CD160 within each concentration; *n* = 3 independent experiments. **c**, HUVEC tube formation at 10 µg mL^−1^ IgG1 or anti-CD160 in unstimulated medium or with VEGF-A (50 ng mL^−1^), bFGF (20 ng mL^−1^) or both. Scale bar, 200 µm. **d**, Quantification of **c** as total tube length: *P* = 0.7268 (unstimulated), 0.1074 (VEGF-A), 0.0311 (bFGF), 0.0300 (VEGF-A plus bFGF). **e**, As **c** for human dermal microvascular endothelial cells (CADMEC/HMVEC), imaged in the green fluorescence channel. Scale bar, 200 µm. **f**, Quantification of **e**: *P* = 0.9987 (unstimulated), 0.1284 (VEGF-A), 0.0027 (bFGF), 0.0115 (VEGF-A plus bFGF). **g**, HUVEC metabolic viability (MTT, % of untreated) at 8, 24 and 48 h with IgG1 or anti-CD160 at 10 µg mL^−1^. **h**, HUVEC viability at 24 h across the antibody dose series (1–20 µg mL^−1^); all IgG1-versus-anti-CD160 comparisons non-significant. **i**, HUVEC lactate dehydrogenase release (% of maximum) at 8, 24 and 48 h at 10 µg mL^−1^. **j–l**, As **g–i** for microvascular endothelial cells. **m**, PeakForce QNM atomic force microscopy topography of HUVEC treated with IgG1 (top) or anti-CD160 (bottom). Scale bars, 1.0 µm (top) and 20 µm (bottom). **n**, Young’s modulus (kPa) extracted from individual force curves. Mann–Whitney test, *P* < 0.0001; three independent experiments, more than 170 force curves per condition, 10 curves per cell.

Because CD160 expression is growth-factor-dependent (Fig. 1), we repeated the assay at 10 µg mL−1 across defined stimulation states in HUVEC (Fig. 2c,d) and in dermal microvascular endothelial cells (Fig. 2e,f). The result mirrors the expression data. In unstimulated cells anti-CD160 had no effect (HUVEC ≈ 28 versus ≈ 28 mm per field, P = 0.7268; microvascular ≈ 24 versus ≈ 24 mm, P = 0.9987). With VEGF-A alone the reduction did not reach significance (HUVEC ≈ −24%, P = 0.1074; microvascular ≈ −13%, P = 0.1284). With bFGF, and with VEGF-A plus bFGF, anti-CD160 reduced total tube length in both cell types (HUVEC ≈ −27%, P = 0.0311 and ≈ −21%, P = 0.0300; microvascular ≈ −29%, P = 0.0027 and ≈ −23%, P = 0.0115). Expanded morphometry reproduces this pattern across network area, mesh, node and segment counts in both cell types (Supplementary Fig. 2b,c). The receptor is functional where it is expressed: the anti-angiogenic response requires the growth-factor conditions that induce CD160.

The phenotype is separable from cell death. Metabolic viability (MTT assay) was flat across antibody dose and time in both cell types, and every IgG1-versus-anti-CD160 comparison at 24 h was non-significant (P = 0.065–0.756; Fig. 2g,h,j,k and Supplementary Fig. 3a). Lactate dehydrogenase release stayed low and indistinguishable between arms (Fig. 2i,l and Supplementary Fig. 3b), with small differences at the highest dose in HUVEC at 8 and 24 h (P = 0.020 and P = 0.032) against an absolute release of ≈ 5–8% of maximum. Suppression of network formation at 10 µg mL−1 therefore occurs with viability and membrane integrity intact, and is distinct from the apoptotic response described for the soluble HLA-G1–CD160 interaction^16^.

Engagement also changed cell mechanics. Atomic force microscopy of antibody-treated HUVEC (Fig. 2m–o) gave a lower Young’s modulus after anti-CD160 than after IgG1 (median ≈ 11 kPa versus ≈ 33 kPa; P = 1.8 × 10−6; Fig. 2n). Cell height rose over the same comparison (median ≈ 3.2 µm versus ≈ 2.3 µm; P = 0.0068; Fig. 2o). A softer, taller, less spread cell is the signature of reduced cortical tension, and it appears at an antibody concentration that leaves viability intact.

### CD160 forms a complex with LRP1 at the endothelial membrane

A GPI-anchored receptor has no cytoplasmic domain, thus it might relay on a co-receptor. We therefore investigated for a transmembrane partner and tested LRP1, on the basis of its role in endothelial proliferation and angiogenesis^31,32,33,34^ and its capacity to couple ligand engagement to autophagy^36^.

Four independent approaches support a CD160–LRP1 complex. Proximity ligation gave a punctate signal for the CD160 + LRP1 antibody pair and essentially no signal with IgG controls, in both HUVEC and microvascular endothelial cells (median ≈ 10–11 puncta per cell versus ≈ 0– 1 in controls; P = 0.0126 and P = 0.0089; Fig. 3d,e). Immunoprecipitation with anti-CD160 from HUVEC lysate (500 µg input) recovered the LRP1 β-chain from the lipid-raft-soluble fraction, with a fainter band in the raft-insoluble fraction and no recovery in the IgG1 lanes (Fig. 3f). Stimulated-emission-depletion (STED) imaging resolved interdigitated CD160 and LRP1 puncta below the confocal limit (Fig. 3g), and confocal imaging showed overlapping LRP1 and CD160 distributions with discrete co-localized objects (Fig. 3h).

**Figure 3.**
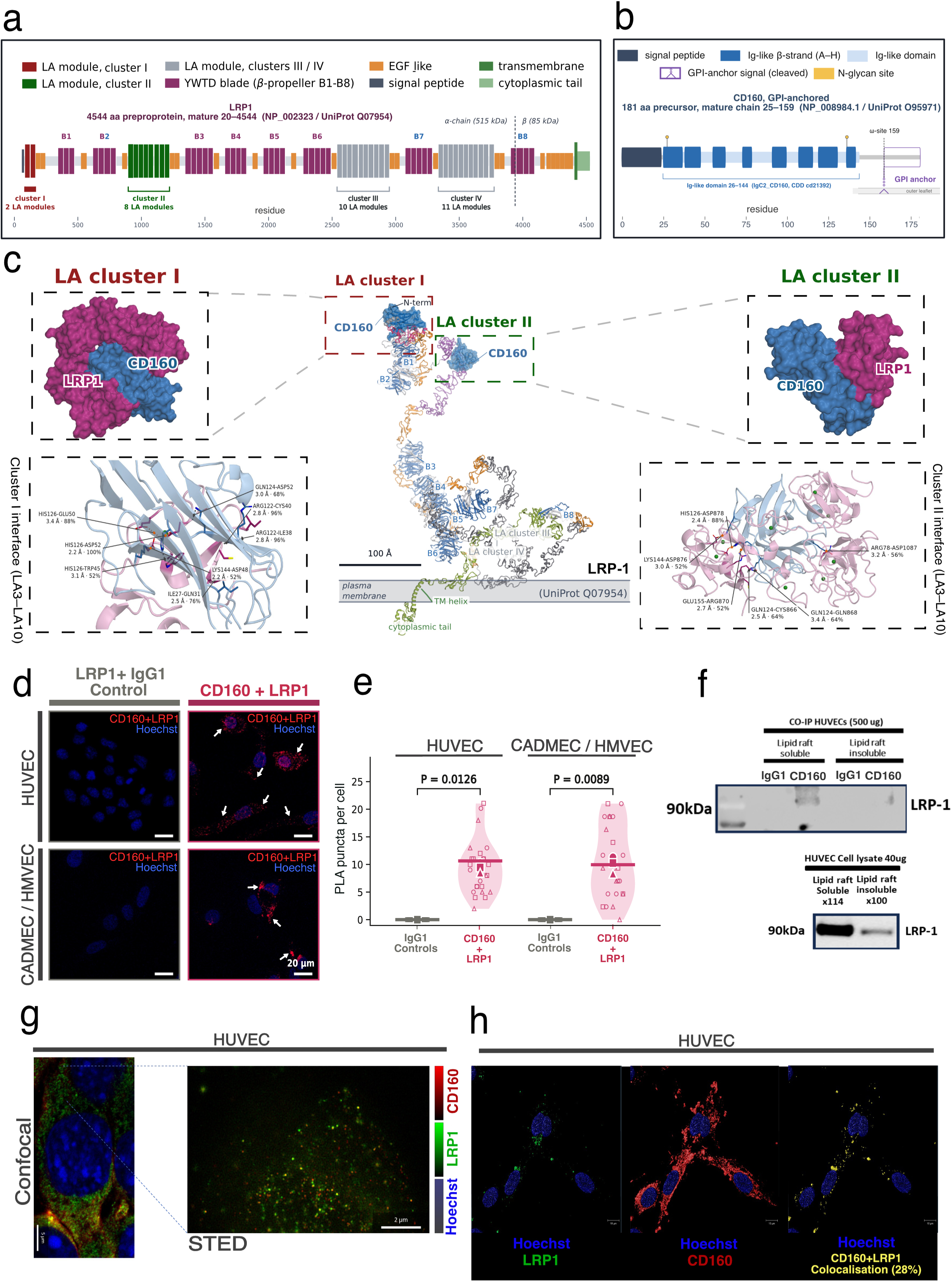
CD160 forms a complex with LRP1 at the endothelial membrane, predicted to engage LA cluster I. **a**, Domain architecture of human LRP1 (UniProt Q07954; 4,544 residues), showing the four clusters of complement-type LA repeats in the α-chain, the EGF and YWTD β-propeller modules, the transmembrane segment and the cytoplasmic NPxY motifs. Clusters I and II are the canonical ligand-binding regions. **b**, Domain architecture of human CD160 (UniProt O95971; 181-residue precursor), showing the signal peptide, the single IgC2-type immunoglobulin-like domain (residues 26–144) and the GPI-anchor ω-site. **c**, Predicted CD160– LRP1 interfaces from Boltz-2 co-folding and ClusPro 2.0 rigid-body docking, rendered as molecular surface with ribbon overlay, with the annotated residue pairs for LA cluster I and LA cluster II. These are computational predictions, not experimental structures; full method comparison, controls and reproducibility statistics are in Supplementary Fig. 4. **d**, Proximity ligation assay in HUVEC (top) and microvascular endothelial cells (bottom) with the anti-CD160 plus anti-LRP1 antibody pair (right) or IgG controls (left). PLA signal in red, Hoechst in blue. **e**, Quantification of **d** as PLA puncta per cell. Welch’s *t*-test versus IgG control: HUVEC *P* = 0.0126; microvascular *P* = 0.0089. *n* = 3 independent experiments with three technical replicates each. **f**, Immunoprecipitation with anti-CD160 or concentration-matched IgG1 from HUVEC lysate (500 µg input) after lipid-raft fractionation, blotted for the LRP1 β-chain. Lanes: raft-soluble and raft-insoluble fractions for each antibody. Representative of three independent experiments.**g**, Confocal and stimulated-emission-depletion (STED) images of CD160 and LRP1 in HUVEC, showing interdigitated puncta below the confocal resolution limit. **h**, Confocal immunofluorescence of LRP1, CD160 and the merge in HUVEC, with co-localised objects. Co-localisation was quantified as Pearson’s correlation coefficient using Coloc2 in Fiji.

We then localized the interface. LRP1 is a 4,544-residue receptor whose α-chain carries four clusters of complement-type LA modules, of which clusters I and II are the canonical ligand-binding regions (Fig. 3a). CD160 is a 181-residue precursor with a single IgC2-type Ig-like domain (residues 26–144) and a GPI anchor at the ω-site (Fig. 3b). We used two independent and mechanistically unrelated computational methods: Boltz-2 co-folding^54^ and ClusPro 2.0 rigid-body docking^55,56^. Both were run across nine LRP1 or LDL-receptor regions, together with a positive control (the RAP D3 domain, whose complex with an LA module is structurally characterized^57^), an off-target arm (LDLR LA3–6) and a binding-site-ablated negative control (Supplementary Fig. 4).

The two methods converge. LA cluster I ranked first of the nine regions on interface predicted TM-score and cleared the 0.6 threshold on every pose, with a median near 0.8 against ≈ 0.45 for the ablated negative control (Supplementary Fig. 4b), and separated from the off-target, ablated and cluster-adjacent arms by Cliff’s δ (nominal P = 0.048–0.095 on seed medians, n = 5 seeds per arm; Supplementary Fig. 4c). The cluster I interface is chemically reproducible rather than a single favorable pose: His126–Asp52 appeared in 25 of 25 poses, Arg122–Cys40 and Arg122– Ile38 in 24 of 25, and His126–Glu50 in 22 of 25 (Supplementary Fig. 4d). Cluster I also won on interface definition rather than interface size: it has a smaller buried surface and a lower interface distance error than the ablated and cluster III arms (Supplementary Fig. 4e). Both methods select the same CD160 face from opposite computational directions. Residues 122, 124 and 144 are hot in both (enrichment P = 1.5 × 10−5; Supplementary Fig. 4f), the top-pose interfaces overlap with a Jaccard index of 0.938 against a random-residue-set background (P = 5.3 × 10−4; Supplementary Fig. 4g), and both light up one contiguous patch centerd on residues 122, 124, 126 and 144. The predicted contact chemistry is the one LRP1 uses generally: a CD160 lysine (Lys144) caps two acidic LRP1 residues adjacent to the module’s calcium site, the recognition logic established for the LRP–RAP complex^57^. The interface residue pairs annotated in Fig. 3c match the cluster I contact frequencies in Supplementary Fig. 4d, and a second set of pairs is annotated for cluster II. These models define a predicted interface, and the complex itself is established by the proximity, fractionation and imaging data above.

### The CD160 anti-angiogenic response requires LRP1

We silenced CD160 or LRP1 in both endothelial cell types (knockdown to ≈ 0.10–0.35 of scramble by RT-qPCR; HUVEC siLRP1 P = 0.0166, siCD160 P = 0.0049; microvascular siLRP1 P = 0.0048, siCD160 P < 0.0001; Supplementary Fig. 1b) and repeated the antibody experiments.

In the scramble background, anti-CD160 reduced total tube length in HUVEC (≈ 42 to ≈ 31 mm per field, ≈ −26%; P = 0.047) and in microvascular cells (≈ 31 to ≈ 25 mm, ≈ −19%; P = 0.039). In both cell types the effect was abolished by silencing CD160 (P = 0.743 and P = 0.359) and by silencing LRP1 (P = 0.805 and P = 0.870) (Fig. 4a–d). The same pattern held for directed migration. Anti-CD160 reduced 9-hour scratch closure in scramble HUVEC (≈ 30% to ≈ 19% of field, a relative reduction of ≈ 37%; P = 0.004) and had no effect in either knockdown (P = 0.728 and P = 0.761), with starting wound areas equivalent across all six arms (P = 0.398–0.612; Fig. 4e–g).

**Figure 4.**
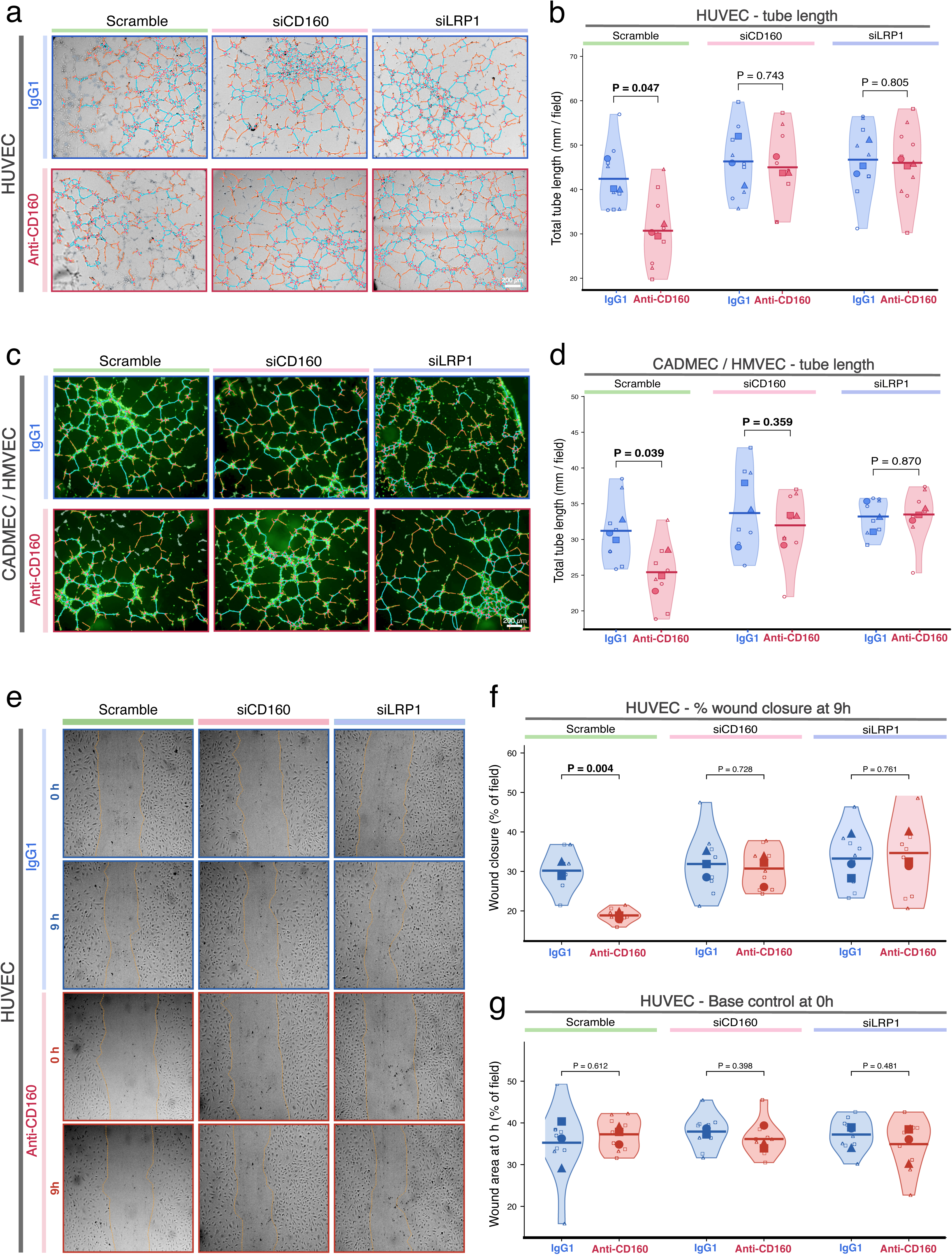
The anti-angiogenic effect of CD160 engagement requires both CD160 and LRP1. **a**, HUVEC tube-formation networks (brightfield with tracing overlay) 48 h after transfection with scramble, si*CD160* or si*LRP1*, treated with IgG1 or anti-CD160 at 10 µg mL^−1^. **b**, Quantification of **a** as total tube length. Welch’s *t*-test, IgG1 versus anti-CD160 within each siRNA background: scramble *P* = 0.047; si*CD160* *P* = 0.743; si*LRP1* *P* = 0.805. *n* = 3 independent experiments. **c**, As **a** for microvascular endothelial cells (green fluorescence channel with tracing overlay). **d**, Quantification of **c**: scramble *P* = 0.039; si*CD160* *P* = 0.359; si*LRP1* *P* = 0.870. **e**, Scratch-wound assay in HUVEC at 0 and 9 h with wound-edge tracing overlay, in the three siRNA backgrounds with IgG1 or anti-CD160. Proliferation was suppressed with mitomycin C. **f**, Quantification of **e** as wound closure at 9 h (% of field): scramble *P* = 0.004; si*CD160* *P* = 0.728; si*LRP1* *P* = 0.761. **g**, Wound area at 0 h for the same arms, confirming equivalent starting geometry (*P* = 0.398–0.612 across all comparisons). Knockdown efficiency for **a–g** is in Supplementary Fig. 1b; viability across the same siRNA and antibody arms is in Supplementary Fig. 5.

Two points follow. The loss of effect in the knockdowns is not a floor effect: under IgG1, tube length in the siCD160 and siLRP1 backgrounds was at or slightly above scramble (≈ 46 versus ≈ 42 mm in HUVEC), so a reduction could have been detected and none occurred. And the antibody effect requires LRP1 as well as CD160, which consolidates the physical association in Fig. 3.. Knockdown itself slightly reduced viability progressively over 48 h (siCD160 and siLRP1 below scramble, P = 0.0069–0.0240 for the cross-condition brackets), yet within every siRNA background the IgG1-versus-anti-CD160 comparison was non-significant at all three timepoints in both cell types (P = 0.2746–0.9515; Supplementary Fig. 5).

### Engagement of CD160 raises autophagic flux through LRP1

LRP1 converts ligand engagement into autophagy^36^, and autophagy controls endothelial sprouting^38,39,40,41,42^. We therefore measured autophagy as flux, using the tandem RFP–GFP–LC3 reporter^52^. Both fluorophores emit in the neutral autophagosome, GFP is quenched on fusion with the acidic lysosome, and the per-cell GFP/RFP ratio therefore falls as autophagosome-to-autolysosome maturation proceeds (Fig. 5a). To separate increased maturation from a maturation block, every arm was run with and without a 10 µM chloroquine clamp^51^, and the whole design was repeated in scramble, siCD160 and siLRP1 backgrounds (Fig. 5b,c).

**Figure 5.**
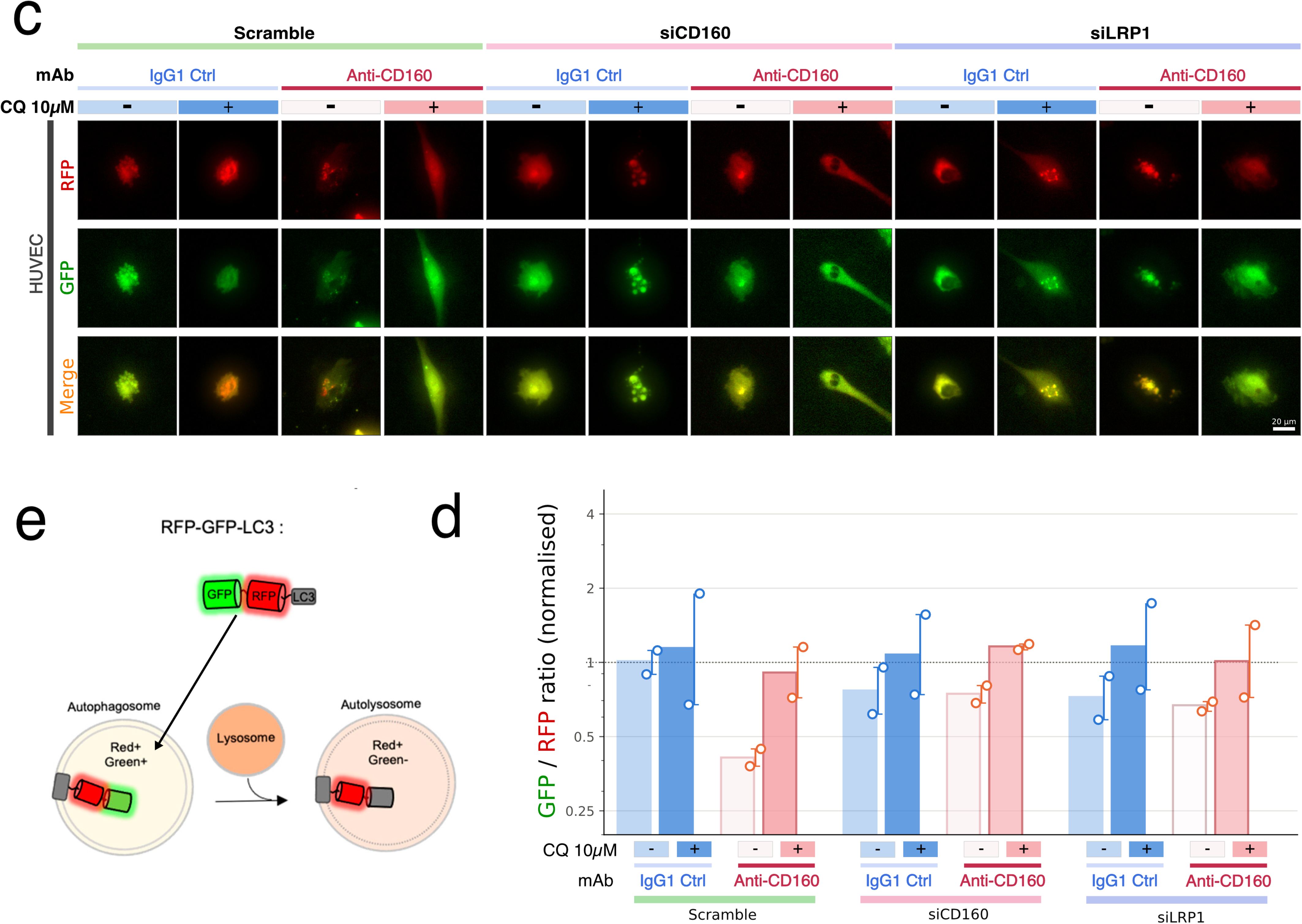
Engagement of CD160 increases autophagic flux, and requires CD160 and LRP1 to do so. **a**, The tandem RFP–GFP–LC3 reporter. Both fluorophores emit in the neutral autophagosome (red-positive, green-positive); GFP is quenched on fusion with the acidic lysosome, so the autolysosome is red-positive, green-negative. A fall in the GFP/RFP ratio therefore reports autophagosome-to-autolysosome maturation, that is, increased flux. **b**, HUVEC expressing RFP–GFP–LC3, transfected with scramble, si*CD160* or si*LRP1*, treated for 24 h with IgG1 or anti-CD160 at 10 µg mL^−1^, each without or with a 10 µM chloroquine clamp for the final 4 h (12 arms). Rows: RFP, GFP, merge. Acquisition settings identical across all arms. Scale bar, 20 µm. **c**, Quantification of **b** as per-cell GFP/RFP intensity ratio, normalised to the scramble/IgG1/vehicle arm (log-scaled axis; dashed line at 1). Anti-CD160 lowered the ratio from ≈ 1.0 to ≈ 0.42 in the scramble background (≈ −58%), an effect absent in si*CD160* (≈ 0.80 versus ≈ 0.78) and si*LRP1* (≈ 0.75 versus ≈ 0.68); chloroquine raised the ratio in every arm, confirming lysosome dependence. At least five cells per field were measured. Error bars, standard deviation.

In the scramble background, anti-CD160 lowered the normalized GFP/RFP ratio from ≈ 1.0 to ≈ 0.42, a reduction of ≈ 58% that corresponds to an increase in flux, and chloroquine returned the ratio to ≈ 0.9. The loss of GFP signal is therefore lysosome-dependent and not a fall in reporter expression. In the siCD160 background the antibody produced no reduction (≈ 0.80 under IgG1 versus ≈ 0.78 under anti-CD160), and the siLRP1 background behaved the same way (≈ 0.75 versus ≈ 0.68). Chloroquine raised the ratio in every arm, as expected for a functioning clamp. The increase in flux driven by receptor engagement therefore requires both CD160 and LRP1, matching the genetic requirement of the anti-angiogenic phenotype in Fig. 4. The IgG1 baseline ratio is lower in both knockdown backgrounds (≈ 0.75–0.80) than in scramble (≈ 1.0), which indicates that silencing either receptor shifts basal flux in its own right.

### Lysosomal function and LDL-receptor-family engagement are required for the anti-sprouting effect

The reporter experiment shows that flux rises when the receptor is engaged. To test whether that flux is required for the phenotype, we blocked the pathway pharmacologically and tested whether the antibody effect survived (Fig. 6). Cells and aortic rings were treated with vehicle, receptor-associated protein (RAP, 500 nM) to block ligand engagement of the LDL-receptor family, chloroquine (10 µM) to block lysosomal degradation, or both, each with IgG1 or anti-CD160. In vehicle, anti-CD160 reduced total tube length in HUVEC (≈ 38 to ≈ 25 mm, ≈ −34%; P = 0.012) and in microvascular cells (≈ 44 to ≈ 27 mm, ≈ −39%; P = 0.004). With RAP the effect was absent (P = 0.581 and P = 0.479). With chloroquine it was no longer detectable (P = 0.187 and P = 0.121), and with both agents it was absent (P = 0.891 and P = 0.827) (Fig. 6a–d).

**Figure 6.**
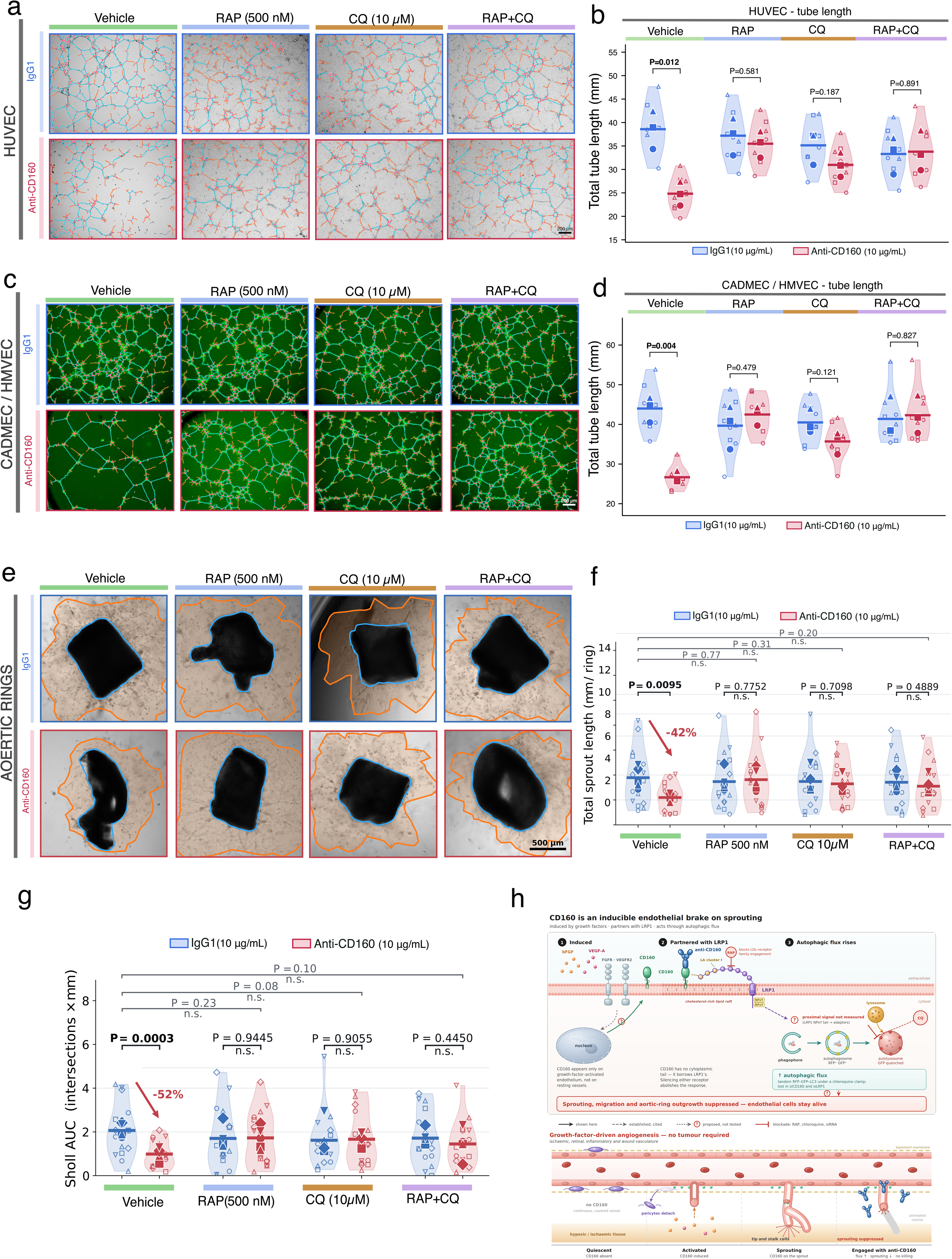
Lysosomal function and LDL-receptor-family engagement are required for the anti-sprouting effect. **a**, HUVEC tube formation with IgG1 or anti-CD160 at 10 µg mL^−1^ under vehicle, RAP (500 nM), chloroquine (10 µM) or RAP plus chloroquine. **b**, Quantification of **a** as total tube length. Welch’s *t*-test, IgG1 versus anti-CD160 within each drug condition: vehicle *P* = 0.012; RAP *P* = 0.581; chloroquine *P* = 0.187; RAP plus chloroquine *P* = 0.891. **c**, As **a** for microvascular endothelial cells. **d**, Quantification of **c**: vehicle *P* = 0.004; RAP *P* = 0.479; chloroquine *P* = 0.121; RAP plus chloroquine *P* = 0.827. **e**, Mouse aortic-ring sprouting at day 7 under the same eight conditions, treatments applied from day 1 and refreshed every 48 h. **f**, Total sprout length per ring. Anti-CD160 reduced sprout length by 42% in vehicle (*P* = 0.0095) and had no effect under RAP (*P* = 0.775), chloroquine (*P* = 0.710) or both (*P* = 0.489). Comparisons across the four drug conditions under IgG1 alone were non-significant (*P* = 0.20–0.77). *n* = 18 animals (three independent experiments of six animals; rings from the same aorta are technical replicates). **g**, Sholl analysis area-under-curve, a measure of sprout arbor complexity, for the same rings. Anti-CD160 reduced Sholl AUC by 52% in vehicle (*P* = 0.0003), with no effect under RAP (*P* = 0.945), chloroquine (*P* = 0.906) or both (*P* = 0.445); IgG1-only comparisons across drug conditions *P* = 0.08–0.23. **h**, Proposed axis. Growth-factor activation induces endothelial CD160; engagement of CD160 in complex with LRP1 increases autophagic flux; flux and LDL-receptor-family engagement are required for suppression of sprouting, with the vessel-level consequences indicated (endothelial cell, basement membrane, pericyte coverage and sprout morphology). Viability and LDH release across all antibody, RAP and chloroquine arms are in Supplementary Fig. 6.

The *ex vivo* aortic ring gave the same response on a multicellular sprouting readout. In vehicle, anti-CD160 reduced total sprout length by 42% (P = 0.0095) and Sholl area-under-curve, a measure of sprout arbor complexity, by 52% (P = 0.0003). Neither readout responded to the antibody in the presence of RAP (P = 0.775 and P = 0.945), chloroquine (P = 0.710 and P = 0.906) or both (P = 0.489 and P = 0.445) (Fig. 6e–g). Under IgG1 alone, comparisons across the four drug conditions were all non-significant (P = 0.20–0.77 for sprout length and P = 0.08–0.23 for Sholl area-under-curve), so RAP and chloroquine did not themselves change baseline sprouting at these doses.

Viability and membrane integrity were monitored across all antibody, RAP and chloroquine combinations at 8, 24 and 48 h in both cell types, using a higher chloroquine dose (25 µM) than the functional assays (Supplementary Fig. 6). The drugs reduced metabolic viability modestly relative to vehicle (HUVEC, 24 h: P = 0.0133 and P = 0.0048 for the cross-condition brackets), yet within each drug condition the IgG1-versus-anti-CD160 comparison was non-significant in vehicle, RAP and chloroquine (P = 0.6444, 0.0722 and 0.7208), with a difference only in the combined RAP plus chloroquine arm (P = 0.0061). The loss of the antibody effect under blockade is therefore not explained by a differential loss of cells between antibody arms.

Together with Fig. 5, these data place autophagic flux and LDL-receptor-family engagement downstream of CD160 and upstream of sprouting suppression. A schematic of the proposed axis growth-factor-induced CD160, antibody engagement, the LRP1 partnership, autophagic flux and suppression of sprouting is given as Fig. 6h.

## Discussion

CD160 has been described on the endothelial cells twenty-five years ago, yet the molecular mechanism of its action has not been fully established. Soluble HLA-G1 was shown to inhibit tube formation, bind endothelial CD160 and trigger apoptosis^16^, and targeting CD160 with an antibody proved anti-angiogenic and vessel-normalizing *in vivo*^17^ and later inhibited ocular neovascularization in rabbit and monkey models^18^. The anti-CD160 antibody we used recognizes both the human and the murine receptor, so the human cell assays and the mouse aortic rings read the same target. The function of CD160 in the endothelium, when it is present, and how its signal reaches the cytoplasm were the open questions. This work adresses them. We show that CD160 is not constitutive in human endothelium, it is induced by bFGF and most strongly by the combination of VEGF-A plus bFGF, and not by TNF-α or TGF-β. Engaging it suppresses network formation, migration, aortic-ring sprouting and arbor network complexity, and softens the cell cortex, while keeping viability and membrane integrity intact. CD160 sits in a complex with LRP1, supported by proximity ligation, raft co-immunoprecipitation, super-resolution imaging and two convergent computational models that nominate LRP1 LA cluster I. Silencing either receptor abolishes the anti-angiogenic response. Engagement raises autophagic flux under a chloroquine clamp, again requiring both receptors, and lysosomal or LDL-receptor-family blockade removes the anti-sprouting effect.

The induction profile CD160 is of a pharmacological interest. The receptor appears under the growth factors that drive pathological angiogenesis, and engaging it suppresses sprouting. Its engagement is therefore the active pharmacology, in contrast to VEGF-pathway inhibition and the same logic that UNC5B^4,5^, DLL4/Notch1^6^ and TIE2 agonism^7^ follow. The functional data is concordant with this, anti-CD160 is inert in unstimulated cells and active in bFGF-containing conditions, which is what an inducible receptor predicts. A therapeutic window arises from this property. A receptor that appears only on growth-factor-activated endothelium is a potential target in tumor, retinal and ischemic vasculature while preserving resting vessels, as opposed to much of the toxicity of VEGF-pathway inhibitors which act on all the endothelium. The vessel-normalizing phenotype of anti-CD160 *in vivo*^17^ and its efficacy in the primate eye^18^ are consistent with an inducible target of this kind.

Both engagements of CD160 described in endothelium suppress angiogenic sprouting, and on that point the present data reproduce the original observation. Soluble HLA-G1 inhibits FGF2-induced tubulogenesis, and the CL1-R2 anti-CD160 monoclonal antibody reproduces that inhibition^16^. The anti-CD160 antibody we used is an agonist, as it engages the receptor and reproduces the direction of the response obtained with the natural ligand soluble HLA-G1. This response is lost when CD160 is silenced, so it is not an off-target effect of the immunoglobulin control, and it is equally lost when LRP1 is silenced, so it runs through the partner that gives this GPI-anchored receptor access to downstream signalling through the cytoplasm. However, the two studies diverge on the fate of the cell. Engagement was reported to induce endothelial apoptosis^16^, whereas at 10 µg mL−1 over 48 h we find metabolic viability and membrane integrity intact, autophagic flux raised and sprouting suppressed.

Three explanations reconcile the two outcomes. First, in the original study the functional readout preceded the death readout: tube formation was scored at 24 h and apoptosis at 50 h, and caspase dependence of the tube phenotype was not tested^16^. The apoptosis reported there therefore cannot be linked directly to the anti-angiogenic effect. Second, Autophagy has already been shown to govern endothelial cell fate. Kringle 5 of plasminogen induces autophagy and apoptotic death in the endothelial cells^44^, and sustained autophagy can precede apoptosis in endothelium exposed to angiogenesis inhibitors^46^. As we demonstrate here, the anti-angiogenic effect is independent of cell death. The intensity, the duration and the ligand (soluble HLA-G1 or an antibody) would then set the rate of autophagic flux and could induce cell death indirectly, as a side effect rather than as the main mechanism behind the CD160 anti-angiogenic effect. Third, the *in vivo* vascular normalizing phenotype described by Chabot et al, following CD160 targeting with CL1-R2 anti-CD160 antibody favors endothelial viability. Vascular normalization is defined by increased pericyte coverage, basement-membrane maturation, reduced permeability and improved perfusion^58,59^ properties that a remodeled, living endothelium acquires. Endothelial apoptosis produces the opposite outcome. It is a component of programmed capillary regression^60^, and withdrawal of endothelial survival signaling selectively ablates immature vessels, from which pericyte-covered vessels are protected^61^. Anti-CD160 raises pericyte coverage and lumen patency^17^ and inhibits ocular neovascularization in two species^18^. A treatment that increases the pericyte coverage which protects vessels from apoptotic regression is nonetheless acting on an endothelium that persists.

LRP1 supplies the cytoplasmic arm that a GPI-anchored receptor lacks^25^, and CD160 has a precedent for association with co-receptors as it can associate with CD2 on NK cells^26^. Molecular docking places CD160 on the LA cluster I surface through the lysine-to-acidic-pair chemistry, adjacent to the module calcium site, that LRP1 uses for RAP and its other ligands^57^. LRP1 has been previously identified as a GPI-protein co-receptor for uPAR^29^, as well as for the cellular prion protein^30^, and has been shown to regulates retinal and tumor angiogenesis^32,33^. Moreover, LRP1 provides the missing link that couples CD160 engagement to autophagy^36,37^. In differentiating chondrocytes, LRP1 loss reduces autophagosome formation, as LC3-II decreases, p62 accumulates and autophagosomes are scarcer by electron microscopy, while the upstream initiators Beclin-1, Fis1 and Perk rise^37^. The upstream signal and the completed flux therefore move in opposite directions in that model. In the endothelial cells studied here, silencing LRP1 shifts basal flux modestly and, more importantly, abolishes the rise in flux that follows receptor engagement. LRP1 is required for the regulated response in both settings, whereas the basal set-point it imposes differs between cell types. The comparison also illustrates why static LC3 levels can point the wrong way, and why every measurement reported here was made under a lysosomal clamp^51^.

The autophagy findings places CD160 in a well-established angioregulatory mechanism. Angiostatic molecules has been shown to consistently involved with endothelial autophagy. For instance, kringle 5 of plasminogen^44^, endostatin through Beclin-1 and β-catenin^45^, decorin through Peg3^47^, endorepellin through angiostasis^48^, and autophagic catabolism of endothelial VEGF-A, which lowers the autocrine drive for sprouting^46,50^. Every member of that series so far is a soluble matrix-derived or proteolytic fragment acting on endothelium.

CD160 extends the series to a cell-surface immunoglobulin-superfamily receptor with a defined transmembrane partner. Its effect aligns with the rest: increased autophagic flux, suppressed sprouting, consistent with the requirement for endothelial ATG5 in ischemia-driven angiogenesis^40^ and with the restraint that PKA-dependent limitation of autophagy places on angiogenesis^42^. The reporter data establish receptor-dependent, lysosome-dependent maturation of LC3 compartments^51,52^. They do not, on their own, establish that the rise in flux is what suppresses sprouting. The pharmacological clamp is consistent with that interpretation but is not decisive, because chloroquine also normalizes tumour vessels through endosomal Notch1 trafficking, independently of autophagy^62^. The decisive test is genetic: silencing ATG5 or ATG7, or expressing a lipidation-dead LC3, and asking whether the anti-sprouting effect of anti-CD160 survives. An interaction-term design across the clamp would quantify that epistasis directly.

Additionally, CD160 modulates endothelial cell mechanical response, as anti-CD160 reduces cell stiffness and spreading, suggesting a link between autophagy and mechanical changes. It lowers the Young’s modulus by about two-thirds and increases cell height, indicating reduced cortical tension. Autophagy can degrade cytoskeletal and focal adhesion components, softening the cell, while a cell with reduced cortical tension sprouts less. The mechanical and the metabolic readouts were taken on the same cells and at the same antibody concentration, and they do not move together: MTT and LDH are flat across the IgG1-versus-anti-CD160 comparison at that concentration and timepoint. The softening is therefore the remodelling of a cell that remains metabolically intact, not the collapse of a dying one, which is the same conclusion the viability data impose on the sprouting phenotype. Stiffness responses are themselves bed-specific, since microvascular endothelial cells from different organs differ in proliferation and migration on the same substrate stiffness^63^.

The clinical potential of targeting CD160 is already supported by strong evidence. CD160 appears on the endothelium of newly formed vessels in human colon carcinoma and mouse B16 melanoma, but not in healthy tissue. Anti-CD160 reduces FGF2-driven corneal neovascularization, oxygen-induced retinopathy, and Matrigel-plug angiogenesis. Combined with chemotherapy, it regresses tumor vasculature and normalizes remaining vessels^17^. The same approach also inhibits ocular neovascularization in rabbits and non-human primates, both alone and in combination with bevacizumab^18^. This work now reveals the mechanism: CD160 is an inducible target acting on the activated endothelium, by inducing autophagy, associating with LRP1. Colorectal cancer highlights the dual potential, as CD160 marks tumor endothelium, while CD160+CD8+ T cells resist exhaustion and restore anti-PD-1 responsiveness via a CD160–PI3K axis. Thus, this same receptor can be targeted in two tumor compartments reverting both neovascularization and T cell exhaustion.^17,64^.

We therefore propose CD160 as an endothelial checkpoint: a receptor that the induced by the activation of the endothelium, whose engagement restrains sprouting. Just as Immune checkpoints became therapeutically druggable once the receptor, its partner and its output were known, and CD160 now meets that standard in the pathological endothelium. Clinical translation of this concept would provide relive to patients who show resistance or toxicity when given VEGF targeted therapies.

## Methods

Extended detail for every method, the full reagent table, image-analysis parameters and per-panel statistics are in Supplementary materials.

### Cells and culture

Pooled primary human umbilical vein endothelial cells^65^ (HUVEC; Lonza, cat. no. C2519A) were expanded in EGM-2 (Lonza). Human dermal microvascular endothelial cells (labeled CADMEC/HMVEC on the figures; PromoCell, cat. no. C-12285, neonatal) were expanded in CADMEC/HMVEC Growth Medium (PromoCell, cat. no. 112-500). Cells were used between passages 3 and 5, were negative for mycoplasma, and were maintained at 37 °C in 5% CO2. Defined stimulation states were established in EBM-2 containing 2.5% serum, to which recombinant human bFGF (20 ng mL−1), VEGF-A165 (50 ng mL−1), both, TNF-α (10 ng mL−1) or TGF-β (5 ng mL−1) was added for 24 h. Each independent experiment used a separate freshly thawed vial of pooled cells, with three technical replicates per condition.

### Antibodies and reagents

Anti-CD160 is a mouse IgG1 monoclonal antibody raised against human and mouse CD160 and produced in-house and provided by professor Benssussan; it is the antibody previously characterized on CD160-expressing tumor cells^66^ and is referred to throughout as anti-CD160. Surface staining of non-permeabilized cells (Supplementary Fig. 1a) places its epitope in the CD160 ectodomain. The isotype control was mouse IgG1 from murine myeloma, clone MOPC-21 (Sigma-Aldrich, M5284), used at the same concentration as anti-CD160 in every experiment. The reference anti-CD160 monoclonal BY55^11^ was used only for the specificity panel in Supplementary Fig. 1a. Other primary antibodies were rabbit monoclonal anti-LRP1 (EPR3724; Abcam, ab92544), rabbit polyclonal anti-VE-cadherin (Cell Signaling Technology, 2158), goat polyclonal anti-PECAM-1/CD31 (M-20; Santa Cruz Biotechnology, sc-1506) for human cells, rat monoclonal anti-mouse CD31 (BD Pharmingen, 557355) for aortic rings, and anti-NG2/CSPG4. Alexa Fluor 488-, 568-and 647-conjugated donkey and goat secondary antibodies (Thermo Fisher Scientific; A21202, A21206, A11001, A11036, A11057, A10037, A21235) were used at 1:2,000. Nuclei were counterstained with Hoechst 33342 (Sigma-Aldrich, B2261) or DAPI. Chloroquine diphosphate (Fluka, 25745), mitomycin C from *Streptomyces caespitosus* (Sigma-Aldrich, M4287), MTT (Sigma-Aldrich, M5655), gelatin from porcine skin (Sigma-Aldrich, G2500), paraformaldehyde (Sigma-Aldrich, P6148), Triton X-100 (Merck), Ponceau S (Sigma-Aldrich, P3504) and hydrogen peroxide (Sigma-Aldrich, H1009; LDH maximum-release control) were used as indicated. Receptor-associated protein (RAP) was used at 500 nM.

Growth-factor-reduced Matrigel was from Corning (354230). A complete reagent table with all catalog numbers, hosts and working dilutions is given as Supplementary Table 1.

### siRNA silencing and RT-qPCR

Cells were transfected with ON-TARGETplus siRNA against human CD160 (Horizon Discovery, L2-019771-01-0010) or human LRP1 (L2-004721-01-0020), or a non-targeting scrambled control, at 25 nM final concentration using DharmaFECT 4 (Horizon Discovery) according to the manufacturer’s protocol, and assayed 48 h after transfection. Total RNA was extracted with the RNeasy kit (Qiagen), reverse-transcribed with the ProtoScript II First Strand cDNA Synthesis kit (New England Biolabs, E6560L), and quantified as 2−ΔΔCt relative to scramble using *RPL32* and *RPS18* as reference genes. Primer sequences are in Supplementary Table 3.

### Flow cytometry

Surface CD160 was detected on the CD160-positive YT line, on CHO cells stably expressing CD160-GPI and on mock-transfected CHO cells, using anti-CD160 or BY55 at 1:300 for 20 min at 4 °C in PBS containing 1% BSA followed by an APC-conjugated secondary antibody, with unstained and secondary-only controls.

### Immunofluorescence, confocal and STED microscopy

Cells were seeded on gelatin-coated glass coverslips, treated as indicated, fixed in 3% paraformaldehyde for 5 min at 37 °C, permeabilized with 0.1% Triton X-100 where intracellular targets were probed, blocked, and incubated overnight at 4 °C with primary antibodies (anti-CD160 1:500; LRP1 1:200; all others 1:500) followed by Alexa Fluor-conjugated secondary antibodies at 1:2,000 for 2 h at room temperature, with Hoechst or DAPI counterstain, and mounted in Fluoroshield without DAPI (Sigma-Aldrich, F6182). Acquisition settings were identical across all arms of an experiment. STED imaging of CD160 and LRP1 used the same staining protocol. Co-localization was quantified as Pearson’s correlation coefficient in Fiji/ImageJ using Coloc2^67,68^. Day-7 aortic-ring outgrowths were stained whole for CD31, CD160 and NG2 by the same protocol.

### Proximity ligation assay

In situ proximity between CD160 and LRP1 was detected by proximity ligation^69^ using the Duolink In Situ Red Starter Kit Mouse/Rabbit (Sigma-Aldrich, DUO92101) according to the manufacturer’s protocol, with the antibody pairs anti-CD160 + anti-LRP1-alpha (Sigma rabit anti-LRP1, L2295) (each 1:500) or mouse IgG1 + anti-LRP1 as the negative control, and quantified as puncta per cell over three independent experiments with three technical replicates each.

### Lipid-raft fractionation, immunoprecipitation and immunoblotting

HUVEC lysate (500 µg protein input) was separated into detergent-resistant and detergent-soluble fractions with the UltraRIPA kit for lipid rafts (BioDynamics Laboratory) according to the manufacturer’s protocol. Immunoprecipitation used anti-CD160 or concentration-matched mouse IgG1. For immunoblotting, whole-cell lysates were prepared in ice-cold RIPA buffer (Thermo Fisher Scientific, 89900) supplemented with 1 mM phenylmethylsulfonyl fluoride, 5 mM sodium fluoride and 1 mM activated sodium orthovanadate, incubated 30 min on ice with vortexing every 10 min, and clarified at 14,000 × g for 15 min at 4 °C. Protein was quantified with the Pierce BCA assay (Thermo Fisher Scientific, 23225); 40 µg per lane was resolved by SDS–PAGE, transferred, probed with the antibodies listed above and HRP-conjugated secondary antibodies, and detected by chemiluminescence. Total protein per lane was measured by Ponceau S staining and used for normalization.

### Angiogenesis assays

Assays follow the consensus guidelines for angiogenesis assays^53^; tube formation is used as a screening assay and the *ex vivo* aortic ring as the multicellular sprouting readout.

Tube formation. Wells of a µ-Slide 15 Well 3D (ibidi, 81506) were coated with growth-factor-reduced Matrigel (Corning, 354230) and allowed to polymerize at 37 °C. Endothelial cells were seeded at 1 × 104 cells per well in 50 µL of the appropriate medium containing the indicated antibody or drug, and networks were imaged after 6 h on an EVOS microscope with a 4× objective, one field per well. Networks were quantified in Fiji/ImageJ with the Angiogenesis Analyzer plugin^70^, reporting total tube length, junction and node counts, segment number, mesh number and network area.

Scratch wound. Confluent HUVEC monolayers were established in a Culture-Insert 2 Well in a µ-Dish 35 mm (ibidi) according to the manufacturer’s protocol; the insert was removed to create the gap, mitomycin C (Sigma-Aldrich, M4287) was added to suppress proliferation, and cells were treated with IgG1 or anti-CD160 and imaged at 0 and 9 h. Closure is expressed as percentage of field, with the 0-hour wound area reported as a control (Fig. 4g).

Mouse aortic ring. Thoracic aortae from 6-week-old male C57BL/6 mice were cleaned of peri-adventitial tissue, cut into 1 mm rings and embedded in growth-factor-reduced Matrigel following the established protocol^71^, then cultured in EGM-2. Treatments were applied from day 1 and refreshed every 48 h; sprouting was scored at day 7 as total sprout length per ring and as Sholl analysis area-under-curve. Experiments were performed and reported in accordance with the ARRIVE 2.0 guidelines^72^.

### Atomic force microscopy

HUVEC were seeded at 5 × 105 per WillCo glass-bottom dish (WillCo Wells, GWST-5040) and left undisturbed for 24 h before treatment. Cells were imaged in PeakForce QNM mode on a Bioscope Catalyst (Bruker) with QNM-LC-CAL probes of nominal spring constant 0.1 N m−1 and tip radius 65 nm, using a 2,000 nm drive amplitude, 0.25 kHz drive frequency and 0.3–0.5 Hz scan rate. Deflection sensitivity was determined by extrapolating the linear portion of three force curves acquired on a non-compliant sample in fluid; the manufacturer’s spring constant was verified by thermal tune in the same medium. Measurements were restricted to regions where cell height exceeded three times the indentation depth to exclude substrate contribution, with 10 curves per cell. Individual force curves were baseline-corrected and the Young’s modulus extracted by contact-point estimation. Cell height was taken from the same topographs. The experiment was performed in triplicate with more than 170 curves per condition, and groups were compared by Mann–Whitney test.

### Viability and cytotoxicity

Metabolic viability was measured by MTT^73^ and membrane rupture by lactate dehydrogenase release (Cytotoxicity Detection Kit LDH, Roche, 11644793001), at 8, 24 and 48 h. Cells were seeded at 2 × 104 per well in 96-well plates in their respective growth medium, treated as indicated, and processed according to the manufacturers’ protocols. Results are expressed as percentage of untreated control (MTT) and as percentage of maximum release (LDH).

### Autophagic flux

Autophagy is reported as flux, in line with community guidelines^51^. HUVEC were expanded in EGM-2 and treated in EBM-2. Cells expressing the tandem mCherry–GFP–LC3B reporter^52^ (pDEST-CMV mCherry-GFP-LC3B WT; Addgene, 123230) were transfected with scramble, si*CD160* or si*LRP1* siRNA, then treated with IgG1 or anti-CD160 at 10 µg mL−1 for 24 h, with chloroquine 10 µM present for the final 4 h where indicated, and fixed and imaged with identical acquisition settings across all twelve arms. The GFP/RFP intensity ratio was measured per cell in at least five cells per field and normalized to the scramble IgG1 vehicle arm. A fall in the ratio indicates autophagosome-to-autolysosome maturation, and its reversal by chloroquine confirms that the change is lysosome-dependent.

### Pharmacological blockade

For the epistasis experiments, cells and aortic rings received vehicle, RAP 500 nM, chloroquine 10 µM, or both, each combined with IgG1 or anti-CD160 at 10 µg mL−1. RAP is an antagonizt of the LDL-receptor family as a whole (LRP1, LRP1B, LRP2, VLDLR, LRP8) and is not LRP1-selective; results obtained with it are reported as family-level blockade.

### Structural modeling

Two independent methods were used, and their outputs are reported as computational predictions rather than experimental structures. Human CD160 (UniProt O95971; RefSeq NP_008984.1) and human LRP1 (UniProt Q07954; RefSeq NP_002323) sequences defined the modeled regions. Co-folding used Boltz-2^54^ with a minimum of five independent seeds per arm, and rigid-body docking used ClusPro 2.0^55,56^. Nine LRP1 or LDL-receptor regions were tested together with the RAP D3 positive control^57^, an LDLR LA3–6 off-target arm and a binding-site-ablated negative control. Poses were ranked by interface predicted TM-score; interfaces were compared by contact frequency across poses, buried surface area, interface distance error, per-residue enrichment and Jaccard overlap against a random-residue-set background.

### Image analysis and statistics

Image analysis was performed in Fiji/ImageJ^67,68^. Within each experiment, acquisition settings, segmentation thresholds and region-of-interest rules were identical across arms and fixed before analysis. For the CD160 and VE-cadherin quantification in Fig. 1b, cell bodies were obtained by nucleus-seeded watershed within a marker-derived foreground defined by minimum cross-entropy thresholding and bounded to 30 µm from the nucleus, and each marker was quantified inside a cell mask derived from the other marker, so that no marker was measured within a mask built by thresholding that same marker. Background was estimated per cell and per channel from a local annulus 1.5–8 µm outside the cell outline, excluding neighboring cells, and subtracted from the cell mean. Cells with more than 2% saturated pixels were excluded from intensity means.

The experimental unit is the independent biological replicate: an independent experiment performed on a separate freshly thawed vial of pooled primary cells, with three technical replicates per condition, or an individual animal for the aortic-ring experiments. Values from individual cells were averaged to a field mean and field means to a replicate mean, and all tests were computed on replicate means (n = 3 independent experiments unless stated otherwise). Graphs show individual replicate and field values together with the replicate means; bars are the mean of replicate means and error bars their standard deviation. Normality was assessed per dataset. Multi-arm comparisons against a single reference condition (Fig. 1b) used one-way ANOVA followed by Dunnett’s multiple-comparison test against the unstimulated arm. Two-group comparisons within a defined condition used Welch’s unequal-variance t-test, with correction for multiple comparisons where several conditions were compared within one experiment. Force-curve distributions (Fig. 2n) were compared by Mann–Whitney test. Analyses were performed in GraphPad Prism and in Python (SciPy). Aortic-ring experiments used 8 animals for Fig. 1c and Fig. 2, and 18 animals for Fig. 6e–g (three independent experiments of six animals, with rings from the same aorta treated as technical replicates). Investigators were not blinded to arm identity.

## Data availability

The data supporting the findings of this study, including source data for every quantified panel, uncropped membranes, image-analysis macros, structural-modeling inputs and output poses, are available from the corresponding authors upon reasonable request.

## Figure legends

**CD160 is an inducible endothelial receptor that suppresses sprouting angiogenesis through LRP1-dependent autophagic flux**

Throughout: the experimental unit is the independent biological replicate (a separate freshly thawed vial of pooled primary cells, with three technical replicates per condition, or an individual animal for aortic-ring experiments). Plots show individual field and replicate values with replicate means; bars are the mean of replicate means and error bars their standard deviation. All tests are computed on replicate means, *n* = 3 independent experiments unless stated otherwise. Marker shapes denote replicate identity (circle, N1; square, N2; triangle, N3). Anti-CD160 and isotype-matched mouse IgG1 were used at the same concentration in every experiment.

## Supplementary figures legends

**Supplementary Figure 1. Antibody specificity and knockdown validation. a**, Flow cytometry of anti-CD160 and the reference BY55 monoclonal on the CD160-positive YT lymphocyte line, on CHO cells stably expressing CD160-GPI and on mock-transfected CHO cells, with unstained and secondary-only controls. Dot plots and histogram overlays. **b**, RT-qPCR validation of knockdown 48 h after transfection, expressed as 2^−ΔΔCt^ relative to scramble with *RPL32* and *RPS18* as reference genes. HUVEC: si*LRP1* *P* = 0.0166; si*CD160* *P* = 0.0049. Microvascular endothelial cells: si*LRP1* *P* = 0.0048; si*CD160* *P* < 0.0001. *n* = 3 independent experiments.

**Supplementary Figure 2. Expanded network morphometry of the tube-formation assays.** IgG1 (blue) and anti-CD160 (red); marker shape denotes replicate identity. **a**, HUVEC antibody dose series (companion to Fig. 2a,b) for network area, node number, segment number and mesh number, with the fitted dose slope per parameter (anti-CD160 versus IgG1 per log_10_ concentration unit: network area −3.66 versus −1.72; nodes −102.3 versus −71.6; segments −42.8 versus −33.7; mesh number is the one parameter with a steeper IgG1 slope). **b**, HUVEC growth-factor-conditioned series (companion to Fig. 2c,d) for the same parameters. **c**, Microvascular endothelial cells, growth-factor-conditioned series (companion to Fig. 2e,f). All *P* values are shown on the panels.

**Supplementary Figure 3. Full viability and cytotoxicity dose–time matrix for antibody treatment. a**, MTT metabolic viability (% of untreated) at 8, 24 and 48 h across the antibody dose series (1, 5, 10, 20 µg mL^−1^) in HUVEC (top row) and microvascular endothelial cells (bottom row). At 24 h, all IgG1-versus-anti-CD160 comparisons were non-significant (*P* = 0.065–0.756). **b**, Lactate dehydrogenase release (% of maximum) for the same matrix. Release remained below ≈ 10% of maximum throughout; the only significant IgG1-versus-anti-CD160 differences were at the highest dose in HUVEC at 8 and 24 h (*P* = 0.020 and *P* = 0.032). *n* = 3 independent experiments.

**Supplementary Figure 4. Boltz-2 and ClusPro 2.0 modelling of the CD160–LRP1 interaction. a**, Interface predicted TM-score for nine LRP1 or LDL-receptor regions, the RAP D3 positive control, the LDLR LA3–6 off-target arm and the binding-site-ablated negative control; points are individual poses with the median marked. LA cluster I ranks first of the nine regions and clears the 0.6 threshold on every pose (median ≈ 0.8 versus ≈ 0.45 for the ablated control). **b**, Separation of cluster I from the off-target, ablated and cluster-adjacent control arms. **c**, Effect sizes (Cliff’s δ) with nominal *P* values on seed medians (*P* = 0.048–0.095; *n* = 5 independent seeds per arm). **d**, Contact frequency across the 25 cluster I poses, labelled as fraction of poses: His126–Asp52, 25/25; Arg122–Cys40 and Arg122–Ile38, 24/25; His126– Glu50, 22/25. **e**, Buried surface area and interface distance error, showing that cluster I wins on interface definition rather than interface size relative to the ablated and cluster III arms. **f**, Per-residue agreement between the two methods, expressed as ClusPro frequency (fraction of 324 models) against Boltz-2 hotness; CD160 residues 122, 124 and 144 are selected by both (enrichment *P* = 1.5 × 10^−5^). **g**, Jaccard overlap of the top-pose interfaces from the two methods against a random-residue-set background (Jaccard 0.938; *P* = 5.3 × 10^−4^). All panels report computational predictions; no experimental structure of a CD160–LRP1 complex is presented.

**Supplementary Figure 5. Viability and cytotoxicity in the siRNA backgrounds.** MTT viability (% of untreated) and LDH release (% of maximum) at 8, 24 and 48 h in scramble, siCD160 and siLRP backgrounds with IgG1 or anti-CD160 at 10 µg mL^−1^, in HUVEC (top) and microvascular endothelial cells (bottom). Knockdown of either receptor reduced viability progressively over 48 h relative to scramble (cross-condition brackets, *P* = 0.0069–0.0240), whereas within every siRNA background the IgG1-versus-anti-CD160 comparison was non-significant at all three timepoints in both cell types (*P* = 0.2746–0.9515). *n* = 3 independent experiments.

**Supplementary Figure 6. Viability and cytotoxicity under RAP and chloroquine blockade.** MTT viability and LDH release at 8, 24 and 48 h in HUVEC (top block) and microvascular endothelial cells (bottom block), for IgG1 versus anti-CD160 at 10 µg mL^−1^ under vehicle (0.1% DMSO), RAP (500 nM), chloroquine (25 µM) or RAP plus chloroquine. Note that this control series uses a higher chloroquine dose than the functional assays in Fig. 6 (10 µM) and is therefore conservative. The drug conditions themselves reduced viability relative to vehicle (HUVEC, 24 h: *P* = 0.0133 and *P* = 0.0048 for the cross-condition brackets), whereas the within-condition IgG1-versus-anti-CD160 comparisons were non-significant in vehicle, RAP and chloroquine (*P* = 0.6444, 0.0722 and 0.7208), with a difference only in the combined arm (*P* = 0.0061). All *P* values are shown on the panels. *n* = 3 independent experiments.

