## Supplementary figures for "CD160 is an inducible endothelial checkpoint that suppresses sprouting angiogenesis through LRP1-dependent autophagy"

### Supplementary Figure 1

a

#### CD160 Antibody validation

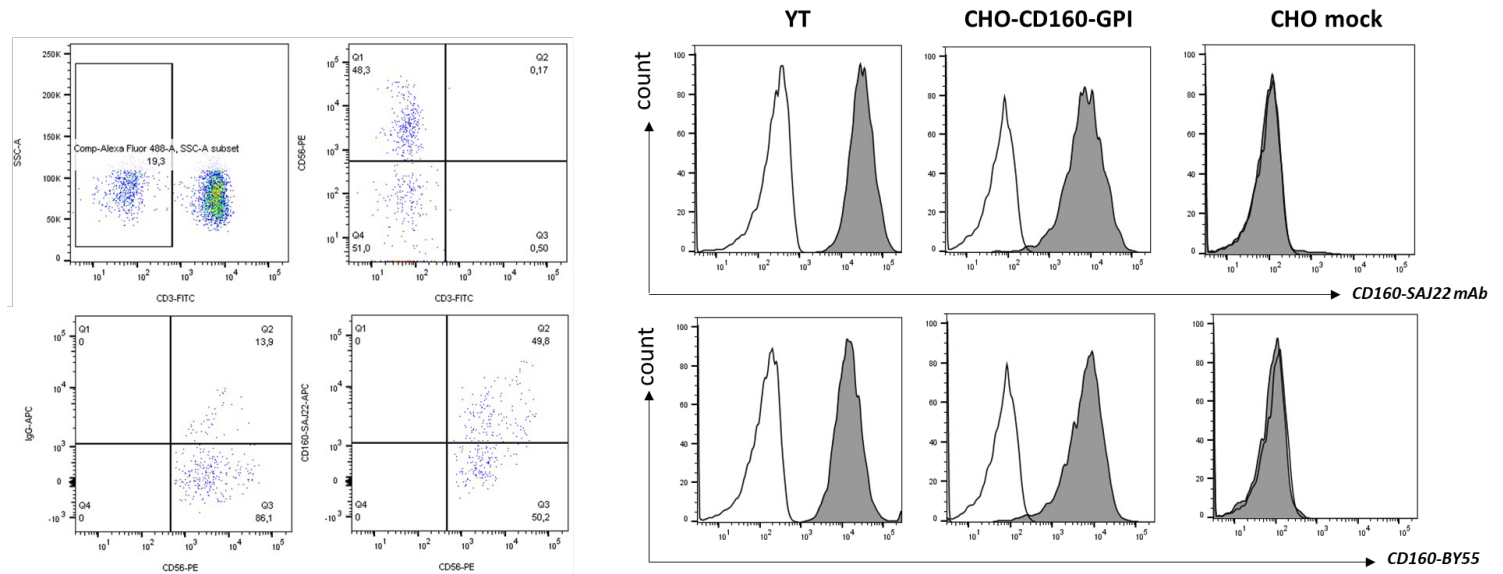

b

#### siRNA knock-downs validation - qPCR

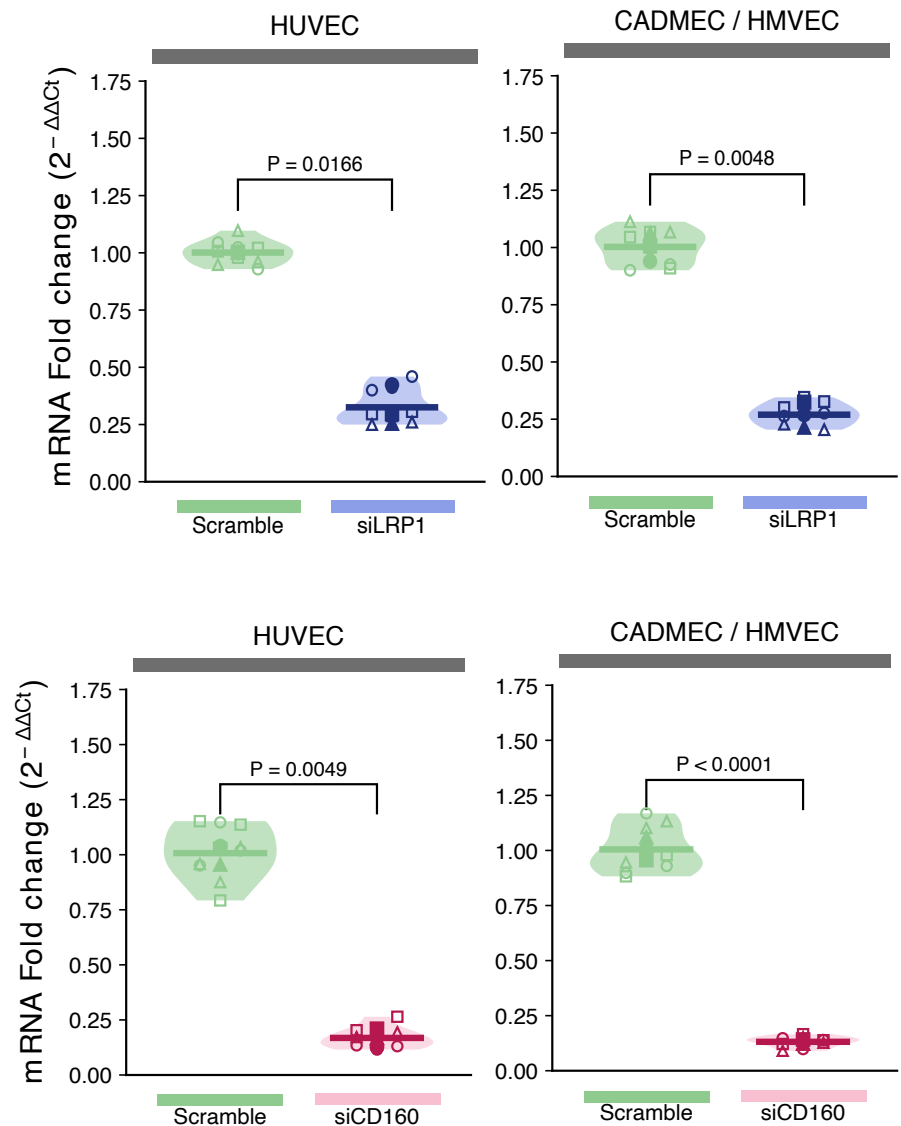

### Supplementary Figure 2.

IgG1 Anti-CD160 ○ replicate N1 □ replicate N2 △ replicate N3 — mean of replicates

#### a HUVEC - Anti-CD160 versus IgG1 control tubogenesis

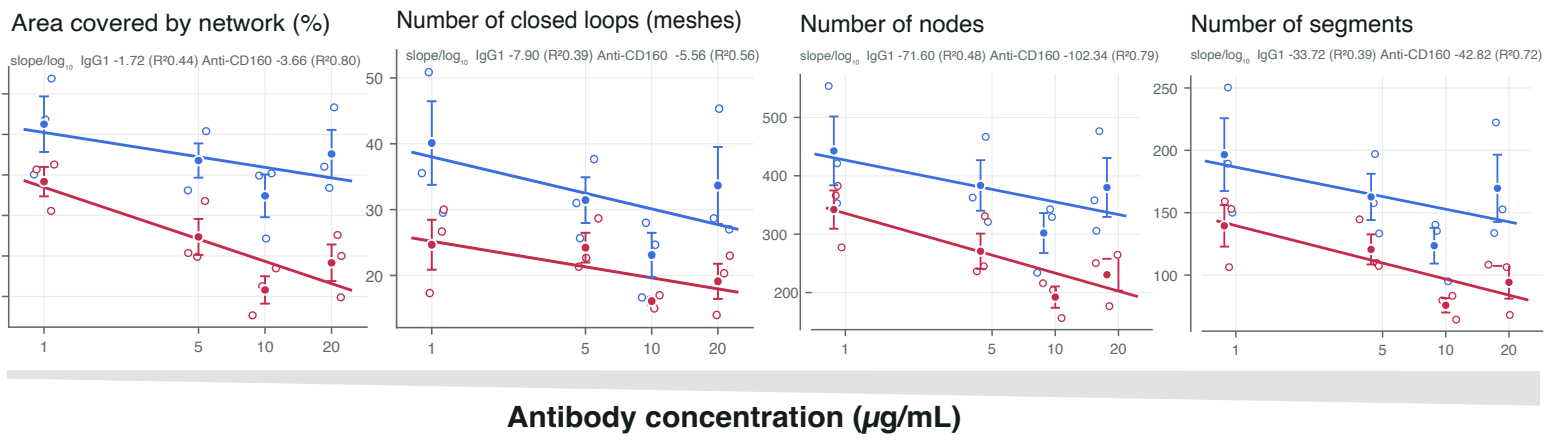

#### b HUVEC - Anti-CD160 versus IgG1 control - Growth factor induced tubogenesis

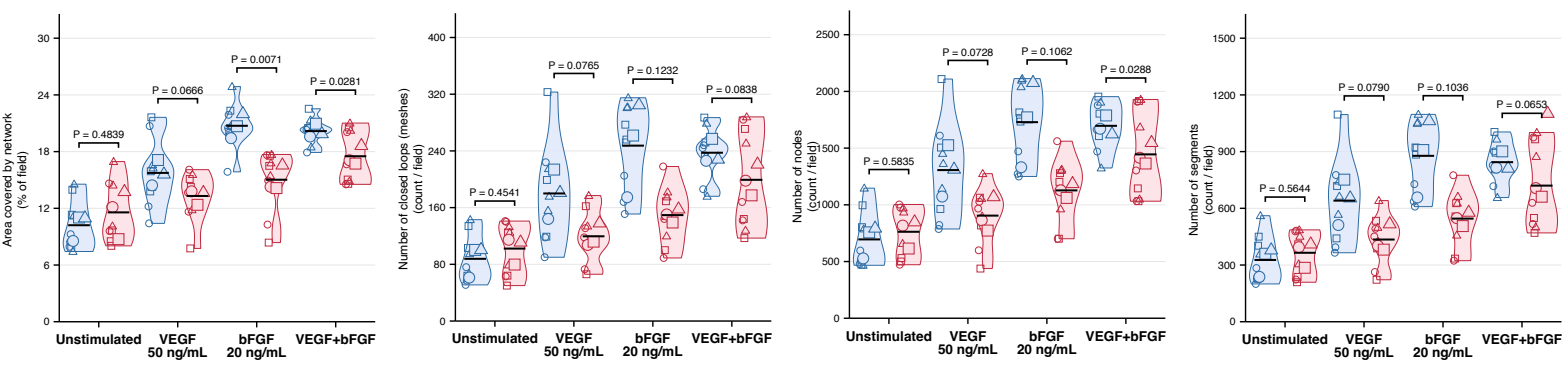

#### c CADMEC / HMVEC - Anti-CD160 versus IgG1 control - Growth factor induced tubogenesis

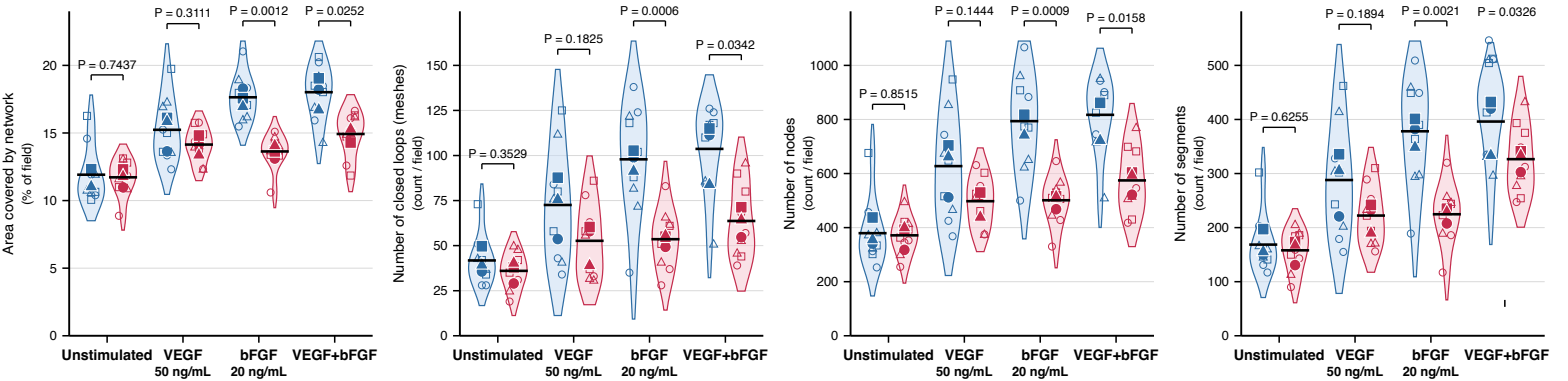

### Supplementary Figure 3

IgG1 SAJ22 replicate N1 replicate N2 replicate N3 — mean of replicates

#### a Anti-CD160 versus IgG1 control - Cell viability (MTT assay)

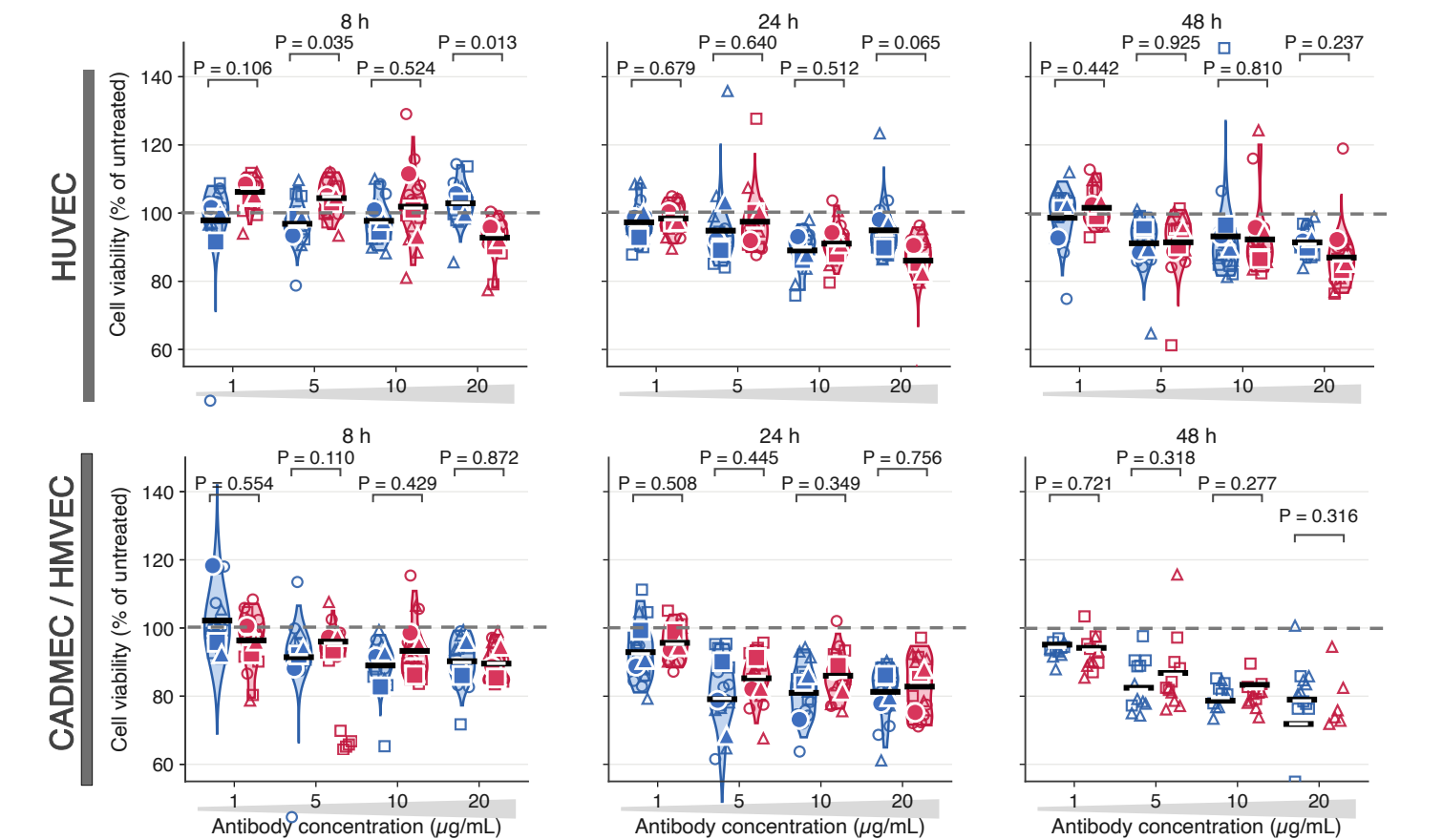

#### b Anti-CD160 versus IgG1 control - Cell death (LDH assay)

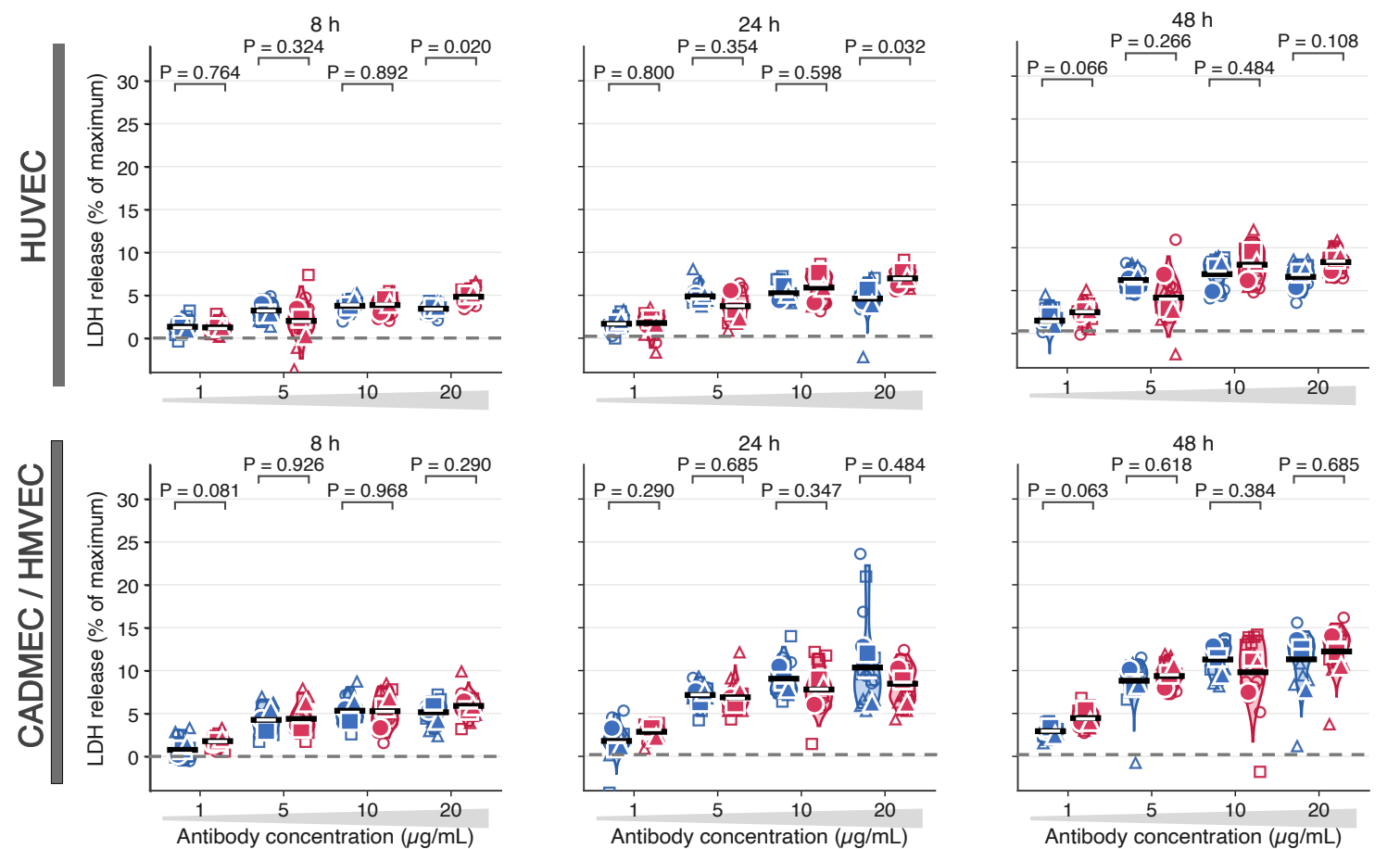

### Supplementary Figure 4

#### Boltz-2 & ClusPro2 modeling of CD160/LRP1 interaction

a

Two independent methods predict that CD160 binds the same patch of LRP1's first ligand-binding cluster

Computational predictions, not experimental structures.

CD160 LRP1 (co-folded) LRP1 (docked) agreed contact residues calcium

The predicted complex: CD160 (blue) docks onto two calcium-binding LRP1 modules (orange); red = Lys144, green = calcium

Close-up: a CD160 lysine caps two acidic LRP1 residues — the grip LRP1 uses for all of its ligands

Boltz-2 co-folding puts the LRP1 module on the red patch

ClusPro 2.0 docking puts the LRP1 module on the red patch

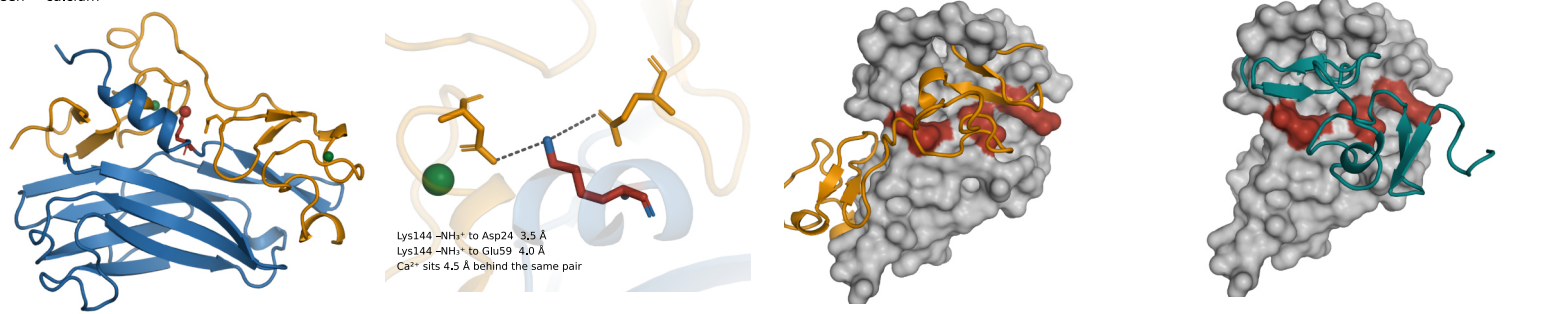

b

Cluster I ranks first of nine and clears the threshold on every pose

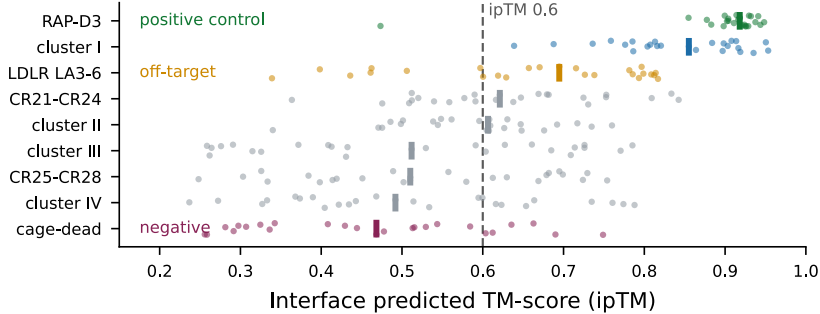

c

Separated from all three controls

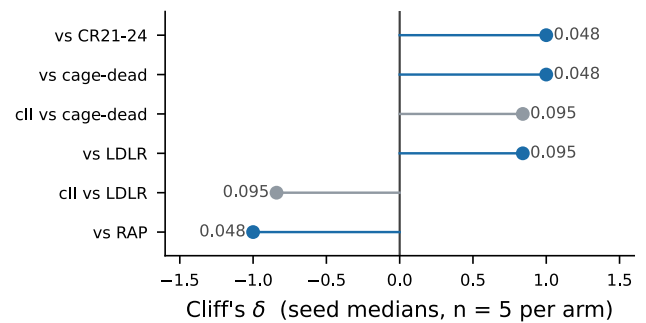

d

The cluster I interface is the same chemistry pose after pose

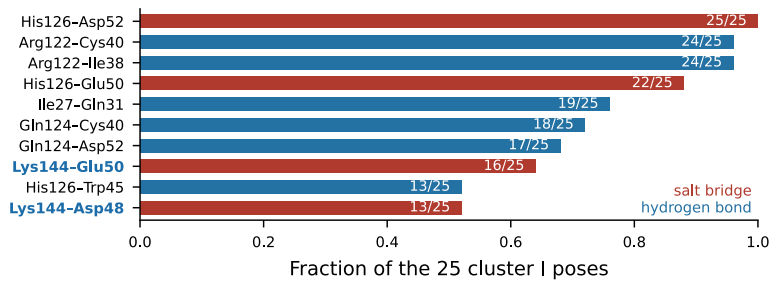

e

Cluster I wins on interface quality, not interface size

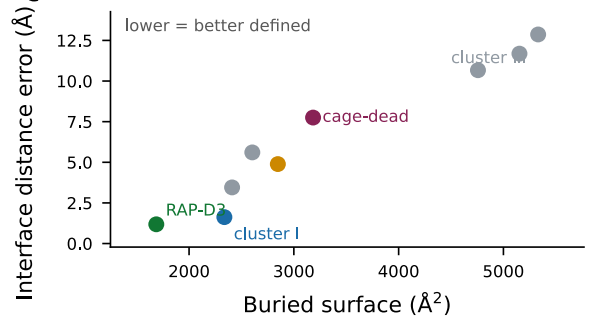

f

Both methods pick the same CD160 hot residues

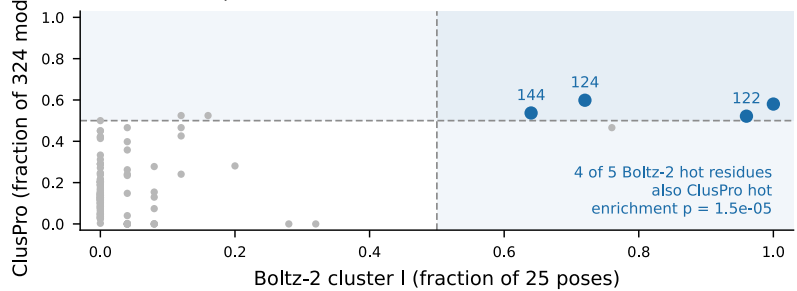

g

Docking recovers the same CD160 face from the reciprocal orientation

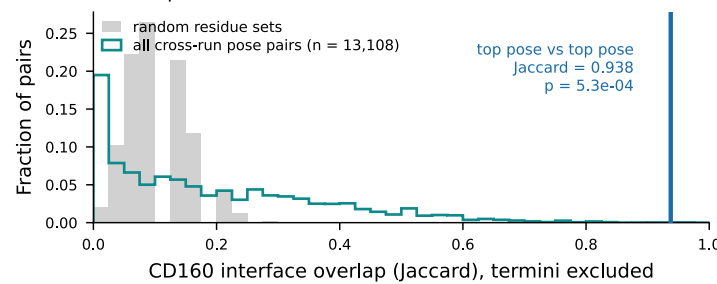

h

Two independent methods light up one contiguous patch

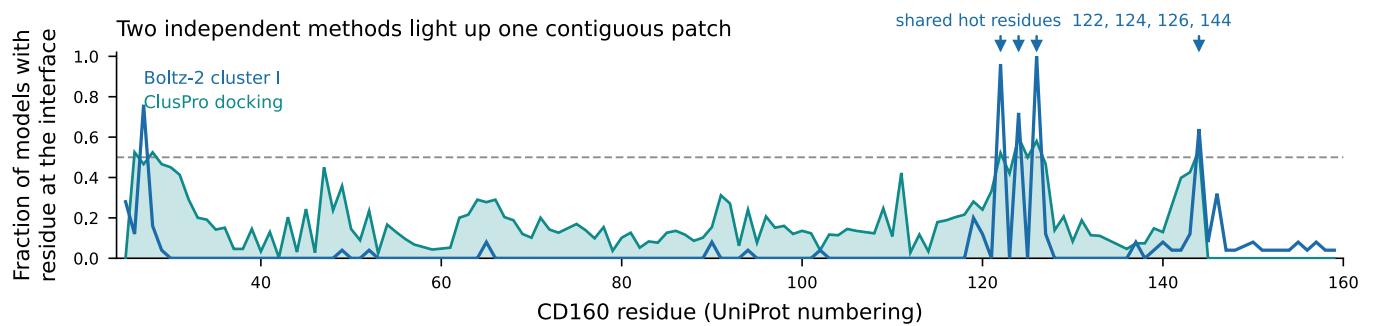

### Supplementary Figure 5

IgG1 Anti-CD160 replicate N1 replicate N2 replicate N3 mean of replicates

a

#### HUVEC - LRP1 and CD160 knock-down -Anti-CD160 versus IgG1 control

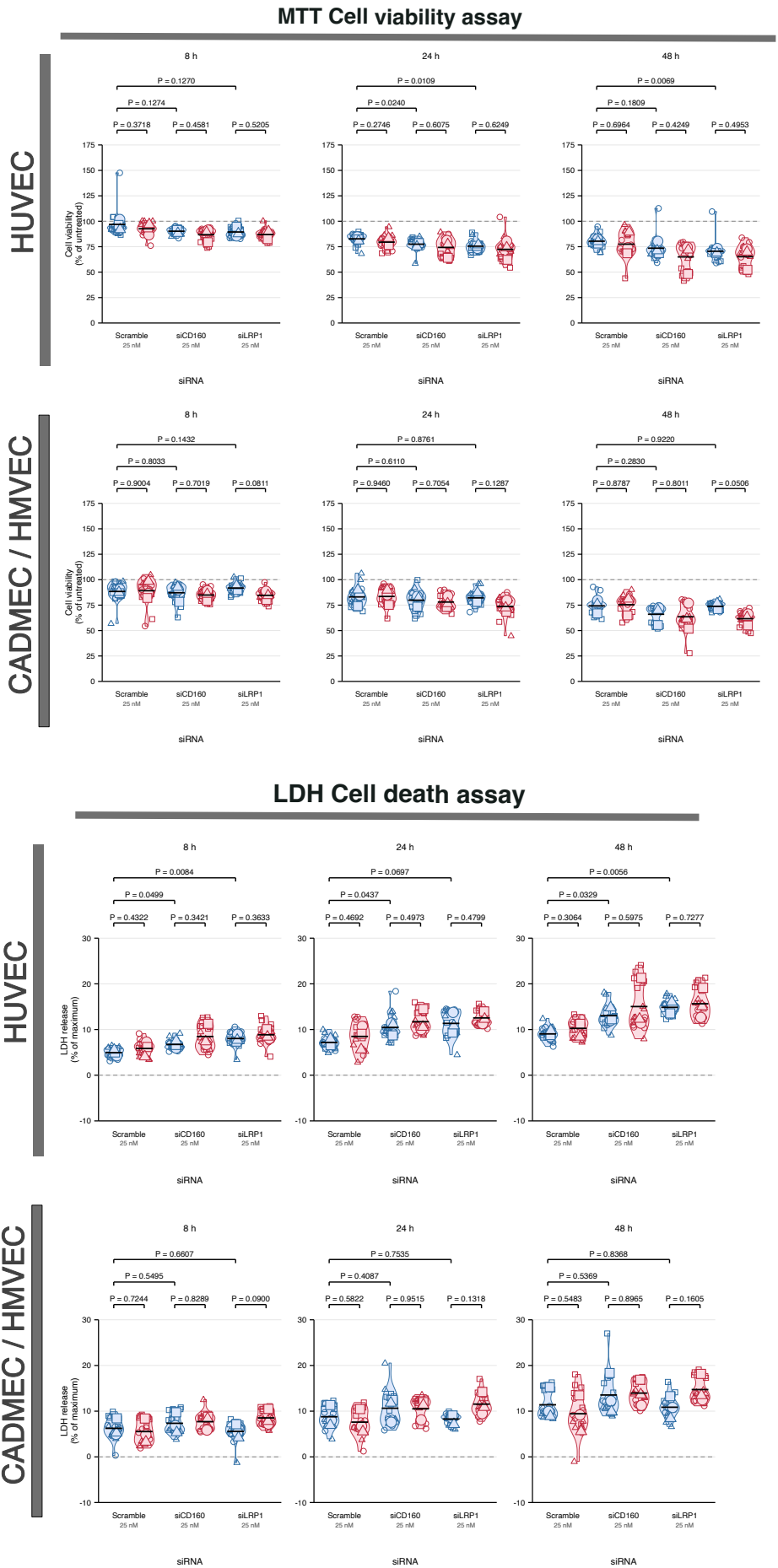

### Supplementary Figure 6

IgG1 Anti-CD160 replicate N1 replicate N2 replicate N3 mean of replicates

#### HUVEC - LRP1 and CD160 +/- RAP at 500 nM and CQ at -Anti-CD160 versus IgG1 control

a

##### MTT Cell viability assay

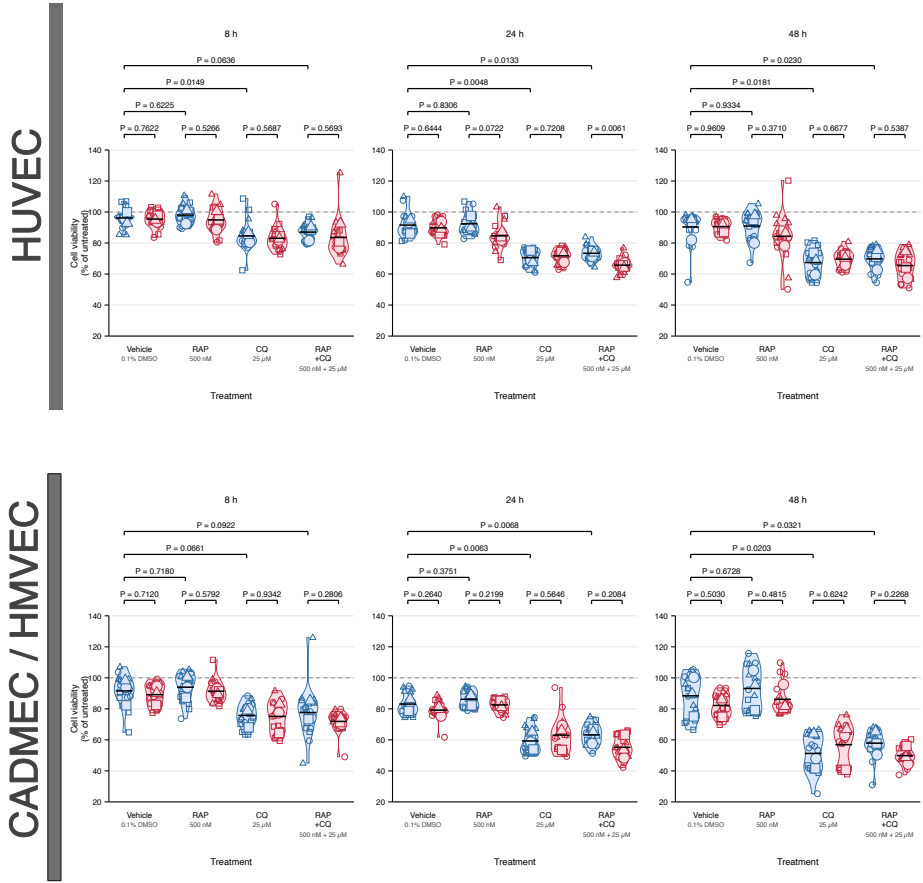

##### LDH Cell death assay

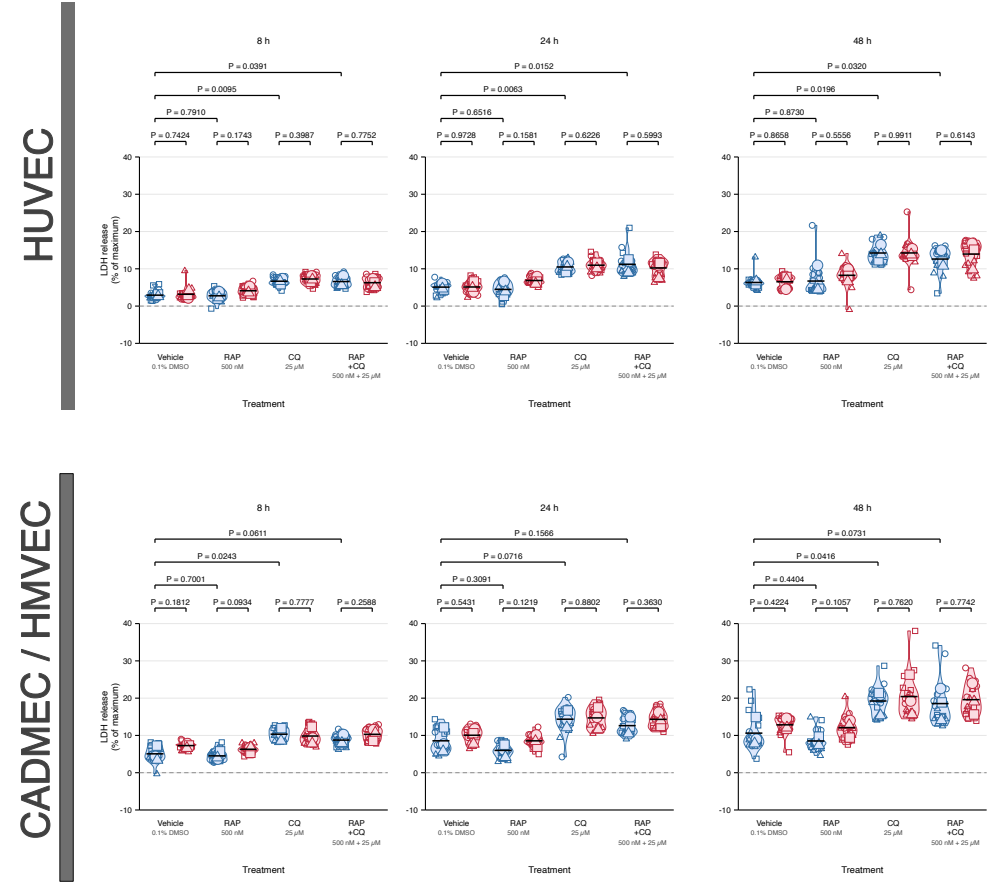
